# Disrupted Inhibitory-Excitatory Balance Underlies Spinal Motoneuron Dysfunction in Incomplete Spinal ord Injury

**DOI:** 10.64898/2026.02.14.705880

**Authors:** Valentin Goreau, François Hug, Louis Simon, Guillaume Le Sant, Raphaël Gross, Thomas Cattagni

## Abstract

Incomplete spinal cord injury disrupts voluntary movement, in part through motoneuron dysfunction, yet the mechanisms underlying this dysfunction remain poorly understood. Using a non-invasive approach to decode the spiking activity of large populations of spinal motoneurons, we quantified the relative contributions of excitatory, inhibitory, and neuromodulatory inputs to motoneuron rate coding after chronic incomplete spinal cord injury. Eighteen participants with incomplete spinal cord injury and 18 age- and sex-matched control participants performed submaximal isometric plantar flexion tasks while high-density surface electromyography was recorded from the soleus and gastrocnemius medialis muscles. Motoneuron firing behaviour was analysed to estimate neuromodulatory drive and the balance between inhibitory and excitatory inputs. Participants with incomplete spinal cord injury exhibited lower rate coding, characterised by lower firing rates at recruitment, lower firing rate modulation, and lower peak firing rates compared with healthy controls. Although estimates of neuromodulatory drive did not differ between groups, individuals with spinal cord injury showed a shift in the inhibition-excitation balance toward greater inhibition compared with controls. Furthermore, increasing inhibitory input through muscle length changes and antagonist tendon vibration modulated motoneuron firing in controls, but not in individuals with incomplete spinal cord injury. Together, these findings suggest that impaired rate coding after incomplete spinal cord injury arises from an altered inhibitory-excitatory balance rather than reduced neuromodulatory drive. Taking advantage of methodological advances to decode spinal motor neuron activity during voluntary contraction, this study identified excessive inhibitory input to spinal motoneurons as a key neural mechanism contributing to muscle weakness and impaired motor function in individuals with incomplete spinal cord injury.

## INTRODUCTION

Incomplete spinal cord injury (SCI) leads to profound alterations in the neural control of movement. Disruption of supraspinal inputs to spinal motoneurons and interneurons at and below the level of the lesion results in muscle weakness and impaired motor function (McDonald & Sadowsky, 2002). Although changes in motoneuron firing characteristics have been described (Thomas et al., 2014), the underlying mechanisms remain poorly understood. In particular, little is known about how SCI alters the relative contributions of excitatory, neuromodulatory, and inhibitory synaptic inputs to motoneuron output.

Muscle force is regulated by modulating the number of active motoneurons (recruitment) and the firing rate of already active motoneurons (rate coding). In individuals with incomplete SCI, a reduction in rate coding has been reported in several muscles (Debenham et al., 2024; Kizyte et al., 2025; Thomas et al., 2014; Wiegner et al., 1993). The mechanisms underlying these alterations remain unclear. Beyond the essential role of excitatory inputs, neuromodulatory inputs contribute to motoneuron rate coding by increasing intrinsic motoneuron excitability through the activation of persistent inward currents (PICs). These currents provide an additional intrinsic depolarising drive that amplifies and prolongs the effects of synaptic excitation in motoneurons (Hultborn et al., 2003; Li & Bennett, 2003; Schwindt & Crill, 1980). Although spinal cord injury can disrupt descending monoaminergic drive, animal studies have demonstrated compensatory mechanisms, such as constitutively active 5-HT_2_ receptors, that preserve PICs despite reduced monoaminergic drive (D’Amico et al., 2014; Murray et al., 2010). Whether PICs are preserved in humans with incomplete SCI remains unknown.

Inhibitory inputs, notably those mediated by Ia interneurons and Renshaw cells, also play an important role in shaping motoneuron output by interacting with both excitatory inputs and PICs (Hyngstrom et al., 2007; Kuo et al., 2003; Powers & Heckman, 2017). Importantly, increases in the net input to motoneurons can arise through distinct patterns of inhibition. One such pattern is the push-pull pattern, in which a decrease in synaptic inhibition occurs in parallel with increased synaptic excitation, thereby supporting higher rate coding and amplifying the effects of PICs (Johnson et al., 2012; Powers & Heckman, 2017; Škarabot et al., 2025). However, after SCI, inhibitory control is altered (Crone, 2003; Knikou & Mummidisetty, 2011), with a notable increase in recurrent inhibition (Shefner et al., 1992). This change may disrupt the normal push-pull interaction between excitation and inhibition and ultimately contribute to firing rate saturation, a phenomenon observed following SCI (Thomas et al., 2014).

Two experimental paradigms are particularly relevant for investigating inhibitory control of motoneuron output. First, changes in muscle length modulate motoneuron excitability through length-dependent inhibitory mechanisms, such as recurrent inhibition (Colard et al., 2025). Specifically, at longer muscle lengths, an increased inhibitory influence is associated with a reduced PIC-mediated prolongation of motoneuron firing rate (Beauchamp et al., 2025; Goreau et al., 2025; Valenčič et al., 2026). Second, vibration applied to the tendons of the antagonist muscles modulates motoneuron output and PIC prolongation effect, via enhanced Ia-mediated reciprocal inhibition (Mesquita et al., 2022; Pearcey et al., 2022). Together, these experimental conditions highlight the flexible and context-dependent regulation of motoneuron output and PIC effects on firing, and therefore provide relevant frameworks for examining inhibitory control of motoneuron output in individuals with incomplete SCI.

In this human study, we aimed to determine how synaptic inputs shape motoneuron rate coding in individuals with incomplete SCI. To this end, we identified large populations of motoneurons (up to 36 per muscle per contraction) by decomposing high-density EMG signals collected during submaximal isometric contractions. We first compared the contribution of inhibitory and neuromodulatory inputs to motoneuron firing rates in plantar flexor muscles between individuals with chronic incomplete SCI and healthy controls, using a framework that links specific motoneuron firing features, such as nonlinear firing acceleration and prolonged firing, to underlying synaptic inputs (Beauchamp et al., 2023; Chardon et al., 2024). We then experimentally manipulated inhibitory inputs through changes in muscle length and antagonist tendon vibration to gain further insights into the role of inhibitory inputs in shaping motoneuron behavior.

## MATERIALS AND METHODS

### Participants and ethical approval

Eighteen individuals with incomplete SCI (mean ± SD : 55 ± 19 years, 175 ± 8 cm, 77 ± 15 kg, two women, Table 1) and 18 control participants (50 ± 20 years, 178 ± 8 cm, 74 ± 14 kg, two women) participated in the study. All participants provided written informed consent, and the experimental procedures were approved by the local ethics committee (CERNI, Nantes Université, IRB: IORG0011023). Participants with SCI had sustained their injury at least five months before enrollment and were classified using the International Standards for Neurological Classification of Spinal Cord Injury (ISNCSCI) as having a C2-L1 lesion. Based on the American Spinal Cord Injury Impairment Scale (AIS), one participant was classified as AIS C, and the remaining 17 as AIS D. Eleven participants had a non-traumatic aetiology. Twelve participants with SCI were taking anti-spastic medication (baclofen and/or pregabalin and/or gabapentin), and five were taking serotonergic medication (duloxetine or paroxetine or mirtazapine) at the time of enrollment. Spasticity was assessed using the modified tardieu scale. Participants with incomplete SCI were required to have had no orthopedic or neurosurgical intervention within the previous three months and no intramuscular injections of botulinum toxin in the plantar flexors during the same period. Control participants were required to have had no lower limb injuries in the previous six months. All participants were instructed not to consume caffeine the day of the evaluation. For individuals with incomplete SCI, the tested leg was the one with the lower maximal voluntary plantar flexion torque, unless this torque was below 15 N.m. In those cases, the limb with greater capacity was selected. As a result, in the incomplete SCI group, 8 of 18 participants were tested on their dominant leg (defined as the leg used to kick a ball), compared with 9 of 18 in the control group.

**Table 1.**
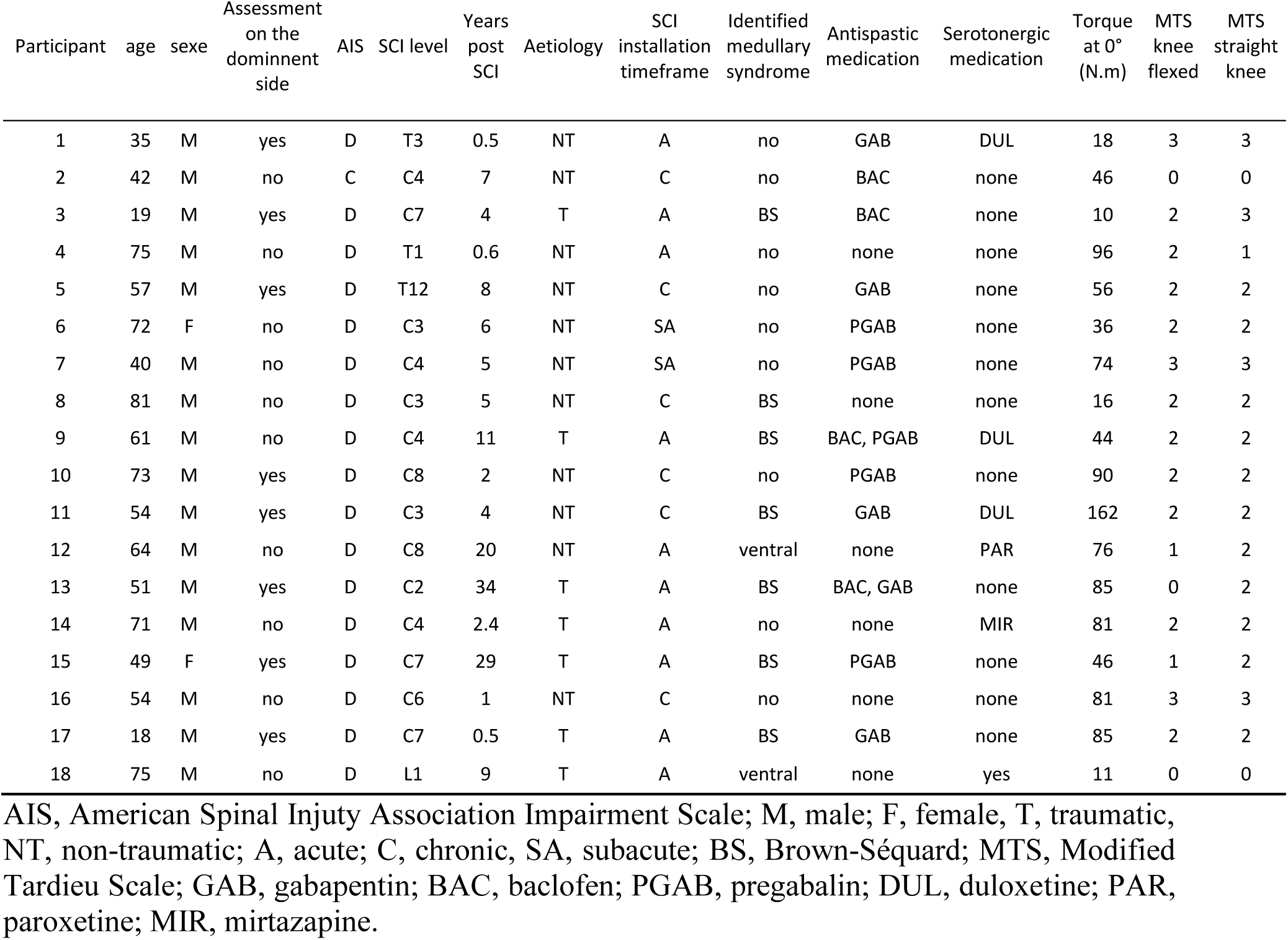
Characteristics of individuals with incomplete spinal cord injury.

The maximal voluntary torque (MVT) of the plantar flexors in the incomplete SCI group was 42.9 ± 33.4 N.m when the muscle was placed at a short length and 62.3 ± 38.0 N.m at an intermediate length (Fig. 1A). In the control group, MVTs were 104.1 ± 33.7 N.m and 145.5 ± 49.6 N.m in the short and intermediate lengths, respectively.

**Figure 1.**
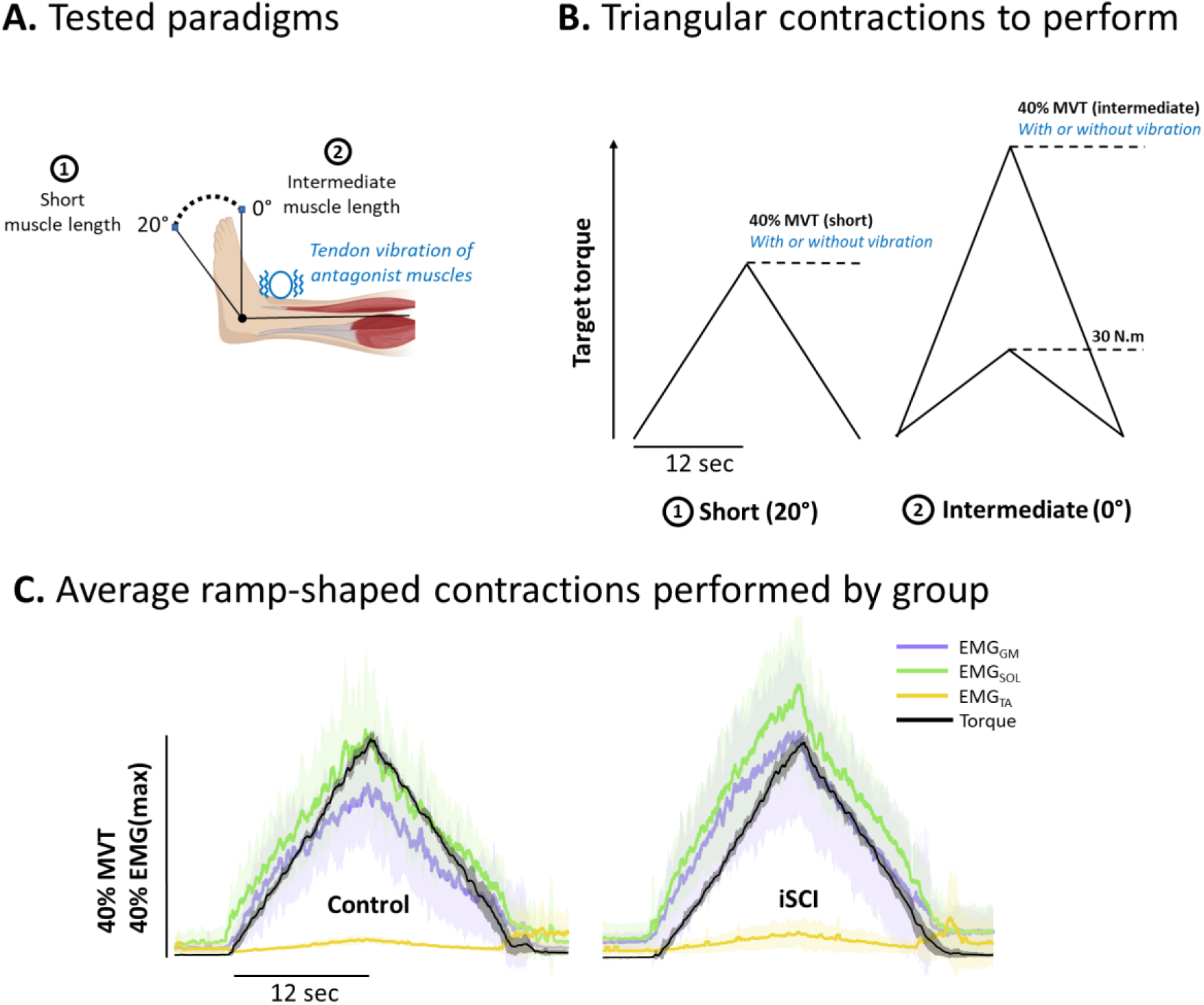
Experimental setup. A, participants performed isometric submaximal plantarflexions at two ankle positions, corresponding to different plantar flexor lengths: 20° (short muscle length position) and 0° (intermediate muscle length position). Tendon vibrations was applied to the antagonist muscles (i.e., dorsiflexor muscles) during contractions to induce reciprocal inhibition. B, the submaximal plantarflexions were performed to 40% of the maximal voluntary contraction torque (MVT) at each position. In addition, a plantarflexion to 30 N.m was performed to compared groups on the same absolute torque. C, mean (thick line) and standard differences (clear outline) of torque and normalised EMG amplitude of gastrocnemius medialis (GM), soleus (SOL) and tibialis anterior (TA), in the control and the incomplete spical cord injury (iSCI group), during the contraction up to 40% MVT in intermediate position.

### Experimental setup

Participants were seated on an isokinetic dynamometer (HUMAC NORM, CMSI, Stoughton, MA, USA), with their foot securely attached to the footplate. Inextensible straps were tightened to immobilise their torso, pelvis, and thigh of the tested leg. The lateral malleolus was aligned with the dynamometer’s axis of rotation. The participant’s position was adjusted to ensure a hip angle of 80° (hip extended = 0°) and a knee angle of 10° (leg extended = 0°). The ankle position was set to either 20° of plantar flexion or 0° (neutral ankle position). Torque signal was digitised at 2048 Hz (Quattrocento, 400-channel EMG amplifier; OT Bioelettronica, Torino, Italia).

### Experimental protocol

Participants began with a standardised warm-up consisting of two 5-second contractions at 50%, 70%, and 90% of their self-estimated maximal contraction intensity, with a 20 s of rest period between contractions. After a 2-minute rest, they performed two 5-second plantarflexion MVTs at each of the two ankle angles, separated by a 2-minute rest period. If the difference between the two contractions exceeded 5%, an additional MVT was performed. Participants received real-time visual feedback of the torque signal during each effort, and strong verbal encouragement were provided. Peak MVT torque was considered as the maximal value obtained from a moving average window of 100 ms and was subsequently used to determine submaximal plantarflexion targets. The same procedure was performed to determine dorsiflexion MVTs.

After the warm-up and MVT assessments, participants performed several isometric submaximal triangular contractions, each consisting of 12-second ascending and descending phases. To compare motoneuron behavior between groups while accounting for large differences in MVT, participants performed a contraction at both a similar relative torque (40% of their MVT) and a similar absolute torque (30 N.m), all performed in the neutral ankle position (0°). To assess how motoneuron behavior adapts to muscle length, participants also performed a contraction at a shorter muscle length (20° of plantarflexion), at 40% of their MVT in this position (Fig 1B). For clarity, these positions are described based on the length of the plantarflexor muscles: the short muscle length position at 20° of plantarflexion and the intermediate muscle length position at 0°. These two joint positions have been shown to induce measurable differences in fascicle length of gastrocnemius medialis (GM) and soleus (SOL) (Colard et al., 2025; Kuzyk et al., 2018).

To assess how reciprocal inhibition affects motoneuron behavior, vibration was applied to the tendons of the antagonist muscles during contractions in both positions (for the contractions at 40% of their MVT). High-frequency vibration (115 Hz; Vibrasens, Technoconcept, Saint-Maurice, France) was applied to the distal tibialis anterior (TA) tendon during plantarflexion contractions by manually pressing the vibrator to the tendon with a force of 15 ± 2 N. Vibration started 10 s before the onset of the triangular contraction and continued until after the torque returned to baseline, ensuring that the entire contraction was covered.

After a familiarisation period for the first condition, participants completed one contraction per condition in randomised order, resulting in a total of five submaximal contractions. At least one minute of rest was provided between contractions to minimise fatigue and the warm-up effect of PICs (Bennett et al., 1998).

### Electromyography

High-density surface electromyograms (EMG) were recorded using two 64 electrode grids per muscle (13 x 5 gold-coated electrodes, with one electrode absent in a corner; 4 mm interelectrode distance; OT Bioelettronica, Torino, Italy) placed over the muscle bellies of both the GM, SOL and TA. The GM and SOL muscles were selected because they are strongly affected by both paresis and signs of hyperexcitability in individuals with incomplete SCI (Jayaraman et al., 2006), whereas the TA was assessed to control for the level of coactivation. The electrode grids were attached using disposable bi-adhesive foam layers (SpesMedica, Battipaglia, Italy), and the cavities in the adhesive were filled with conductive paste (SpesMedica, Battipaglia, Italy). A reference electrode (Kendall Medi-Trace, Canada) was placed over the patella, and a ground electrode (strap dampened with water) was placed around the ankle of the tested leg. The EMG signals were recorded in monopolar mode, bandpass filtered (10–500 Hz), and digitised at a sampling rate of 2048 Hz (EMG-Quattrocento, 400 channel EMG amplifier; OT Bioelettronica, Torino, Italy). The data were acquired using OT BioLab+ software (OT Bioelettronica, Torino, Italy). To obtain information on muscle activation, we reproduced a bipolar configuration by averaging two groups of 25 channels located at the extremities of each grid and then computing the difference between these averages, yielding a single value per grid. The values from the two grids were subsequently averaged to obtain a single value per muscle. The EMG amplitude was analysed as a normalised root mean square over a 300 ms window, by normalising RMS values to the RMS value of the same electrode during the maximal voluntary contraction (EMGmax).

### Decomposition of electromyography signals

We used the convolutive blind source separation method (Negro et al., 2016) implemented in an open-source software (Muedit; Avrillon et al., 2024) to decompose the EMG signals into motor unit spiking activity. The automatic decomposition was performed on the monopolar EMG signals (noisy channels excluded) from one contraction per task and condition. Then, we manually edited all the motor unit spike trains following previously published procedures (Del Vecchio et al., 2020; Hug et al., 2021). Importantly, this procedure has demonstrated high reliability across operators (Hug et al., 2021). We manually edited all motor units in each triangular contraction. The editing process involved an iterative approach, discarding detected peaks that result in erroneous firing rates (outliers) and adding missed firing times that are clearly distinguishable from the noise. The motor unit pulse trains were recalculated with updated motor unit filters and were accepted by the operator once all the putative firing times were selected. For more details on the manual editing step, please see Avrillon et al. (2024).

### Motor unit action potentials amplitude

Motor unit action potentials (MUAPs) were extracted using spike-triggered averaging of high-density surface EMG signals. Single differential EMG signals were computed along the electrode columns and used for subsequent analysis. Spike-triggered averaging was performed using all the firing times identified by decomposition as triggers. MUAPs were extracted using a 170-sample window centered on each firing time (∼12 ms), and the resulting averaged waveforms were detrended. For each motor unit, peak-to-peak amplitude was calculated for each available single differential channel as the difference between the maximum and minimum values of the averaged MUAP waveform. The MUAP amplitude for a given motor unit was defined as the maximum peak-to-peak value observed across all channels. This procedure was applied to all identified motor units, and MUAP amplitude values were subsequently grouped according to the corresponding muscle.

### Motoneuron firing rate

Recruitment threshold of each motoneuron was defined as the torque at its first firing. Instantaneous firing rates (in pulspe per second; pps) were then smoothed using support vector regression to create continuous estimates (Beauchamp et al., 2022). From these smooth profiles, the firing rate at recruitment was taken as the first value, and the peak firing rate was taken as the maximal value. Firing-rate modulation was calculated as the difference between peak firing rate and firing rate at recruitment.

Using a reverse-engineering approach, previous studies have shown that the contribution of inhibitory and neuromodulatory inputs to the motoneuron behavior can be inferred from specific features of the firing rate (Beauchamp et al., 2023; Chardon et al., 2024). Specifically, non-linearities in motoneuron firing during the ascending phase of triangular contractions were quantified using a geometric approach (Beauchamp et al., 2023). The geometric approach assesses how much the motoneuron firing rate deviated from a linear relationship with time output during the ramp-up phase of the contractions (Figure 2B). In this approach, the smoothed firing rate of each motoneuron was plotted against the time generated during a contraction. The largest perpendicular distance between this firing rate and a hypothetical linear increase from recruitment to peak firing rate was quantified, and this distance was defined as brace height. To ensure that brace height reflects only the relative deviation from a linear pattern and to eliminate any bias from differences in peak firing rates across groups, brace height was normalised to the height of a right triangle, where the hypotenuse runs from the point of recruitment to the peak firing rate. We excluded firing trains of motoneurons if they showed a negative slope during the acceleration phase or had a negative brace height before normalisation (567/5222 for control; 725/5113 for incomplete SCI). This method splits the firing rate into acceleration and attenuation phases, which could unintentionally create an artificial attenuation phase if the true attenuation phase had not been reached during the ascending phase of the triangular contraction. To avoid this, motoneuron firing trains were excluded if the duration between recruitment and peak firing rate was shorter than 3 s (1529/5222 for control; 1454/5113 for SCI). In addition to brace height, we calculated the slopes of both the initial acceleration phase and the subsequent attenuation phase of the motoneuron firing rate (Fig 2). Modeling studies (Beauchamp et al., 2023; Chardon et al., 2024) have shown that (i) *brace height* is specifically sensitive to the level of neuromodulation, which is likely proportional to monoamine release and enhances the effective strength of PIC channels; (ii) *attenuation slope* is specifically sensitive to the pattern of inhibition, that is how synaptic inhibition covary with synaptic excitation, such that a steeper attenuation slope reflects a stronger push-pull inhibitory pattern, meaning inhibition decreases as excitation increases; and (iii) *acceleration slope*, reflecting the amplification effect of PICs on firing rate (Avrillon et al., 2024; Revill & Fuglevand, 2017), is the result of the combined influence of neuromodulatory inputs and of the pattern of inhibition.

**Figure 2.**
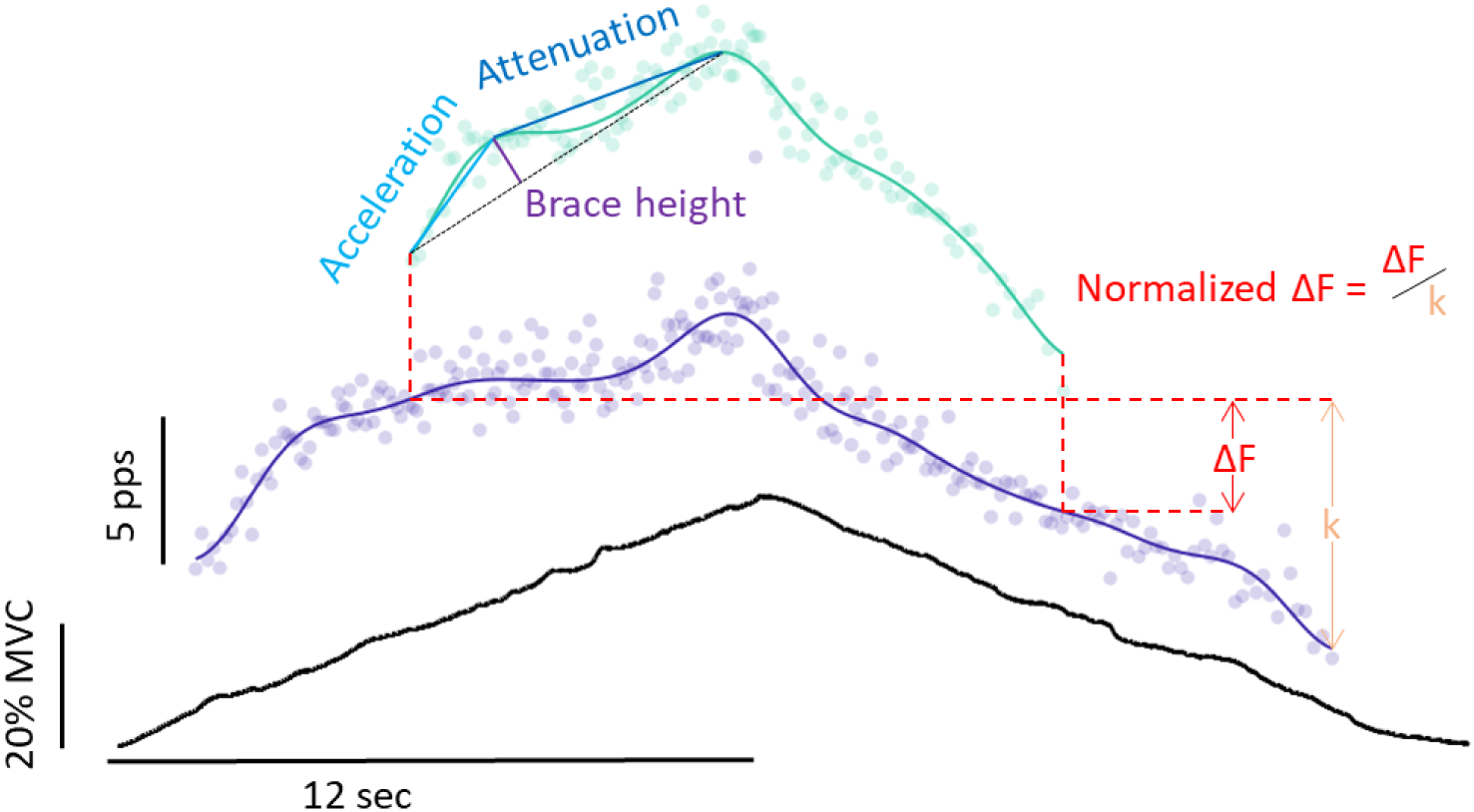
Analyses from motoneuron firing rate. The geometric approach was used to calculate brace height, acceleration, and attenuation, which were used to estimate the level of neuromodulation, PICs amplification effect on the firing, and the pattern of inhibition, respectively. Persistent inward currents (PICs) prolongation effect on firing were estimated using the paired motoneuron analysis (ΔF). In this example, the test motoneuron’s smoothed firing rate is shown in green, while the control units’ smoothed firing rates are shown in purple. Normalised ΔF is calculated by dividing ΔF by the difference in the control motoneuron at test motoneuron recruitment and the firing rate at control motoneuron derecruitment.

Non-linearities in motoneuron firing during the descending phase of triangular contractions were quantified using the paired motoneurons analysis (Gorassini et al., 2002). For this analysis, motoneurons with a lower recruitment threshold (control units) were paired with motoneurons with a higher recruitment threshold (test units). The difference in the firing rate of the control motoneuron between the time at recruitment and the time at derecruitment of the test unit was calculated and referred to as ΔF, as shown in Fig. 2A. In agreement with previous work, the criteria for including a pair of motoneurons were as follows: 1) the test motoneuron should firing for at least 2 seconds, 2) the test motoneuron should be recruited at least 1 second after the control motoneuron to ensure full activation of PIC, 3) the test motoneuron should be derecruited at least 1.5 seconds before the control motoneuron to avoid overestimation of ΔF, and 4) a coefficient of determination (r²) > 0.7 should be observed between the smoothed firing rates of the test and control motoneuron (Hassan et al., 2020). ΔF values are presented as "unit-wise" averages for all suitable test-control motoneuron pairs, thus reducing the number of ΔF values to one value per test unit. *ΔF* reflects the prolongation effect of PICs on firing rate and is the result of the combined influence of neuromodulatory inputs and of the pattern of inhibition (Beauchamp et al., 2023; Chardon et al., 2024). Because ΔF depends on the firing rates of the control unit, which differ between the two populations studied here, we complemented the analysis by normalising ΔF to the maximal value that could have occurred based on the modulation of the control motoneuron. Specifically, ΔF was divided by the difference in the control motoneuron firing rate at test motoneuron recruitment and at control motoneuron derecruitment (i.e., “k”, Fig. 2) (Škarabot et al., 2025).

### Statistical analyses

To assess differences in motoneuron firing characteristics between groups during contractions at 40% MVT and 30 N.m performed at an intermediate muscle length, we applied a linear mixed-effects model to each motoneuron firing metric (i.e., recruitment threshold, firing rate at recruitment, peak firing rate, firing rate modulation, ΔF, normalised ΔF, brace height, acceleration, attenuation and MUAP amplitude). The models included the group (incomplete SCI, control), muscle (gastrocnemius medialis, soleus) and their interactions as fixed effects. The full random-effects structure consisted of a random intercept for each participant and random slopes to account for between participants variability in the fixed effects [e.g., lmer(metric ∼ group*muscle + (group*muscle|id_participant))]. For each model, the random-effects structure was selected by minimising the bayesian information criterion across all plausible random-slope structures. The random slope for the primary effect of interest (group) was always retained, as it represents the key source of between-participant variability for the tested condition.

To test the effects of position and vibration, separate models were fitted including the effect of interest (position: short, intermediate; or vibration: on, off), together with muscle (gastrocnemius medialis, soleus), group (incomplete SCI, control), and their interactions as fixed effects. The random-effects structure was selected using the same procedure; however, in these models the retained random slope corresponded to the effect of interest (position or vibration), rather than group.

To compare TA coactivation between groups, we compared TA EMG amplitude during the similar relative torque contraction at the intermediate muscle length (shown in Fig. 1C) using a t-test. We considered the EMG normalised to the maximal value obtained during the maximal dorsiflexion contraction.

All analyses were performed in R (v.4.4.0; R Core Team 2021, R Foundation for Statistical Computing, Vienna, Austria). Statistical significance was assessed using the lmerTest package in R (Kuznetsova et al., 2017), which uses Satterthwaite’s type III method to approximate degrees of freedom and generate p-values for mixed effects models. Model assumptions of homoscedasticity, normality, and independence of residuals were verified graphically. When significant effects were detected, Sidak post-hoc corrections were applied. Estimated marginal mean differences with 95% confidence intervals and standardised effect sizes (Cohen’s d) were calculated using the emmeans package. The alpha level for all statistical tests was set at 0.05.

## RESULTS

### Motoneuron yield and TA coactivation

In the incomplete SCI group, an average of 9.3 ± 7.5 motoneurons for the GM and 10.3 ± 8.1 motoneurons for the SOL were identified per contraction and participant. In the control group, an average of 13.5 ± 7.8 motoneurons for the GM and 8.2 ± 6.1 motoneurons for the SOL were identified per contraction and participant. TA coactivation during triangular contractions did not differ between groups when assessed using normalised EMG (control vs. incomplete SCI: 2.80 ± 4.07 vs. 3.13 ± 1.68 % of EMGmax, p = 0.7965).

### Differences between groups at intermediate muscle length (0°)

#### Recruitment threshold, rate coding and motor unit action potential amplitude

An example of smoothed motoneuron firing rate from a single contraction in one control and one incomplete SCI participant (#10) is shown in Figure 3.

**Figure 3.**
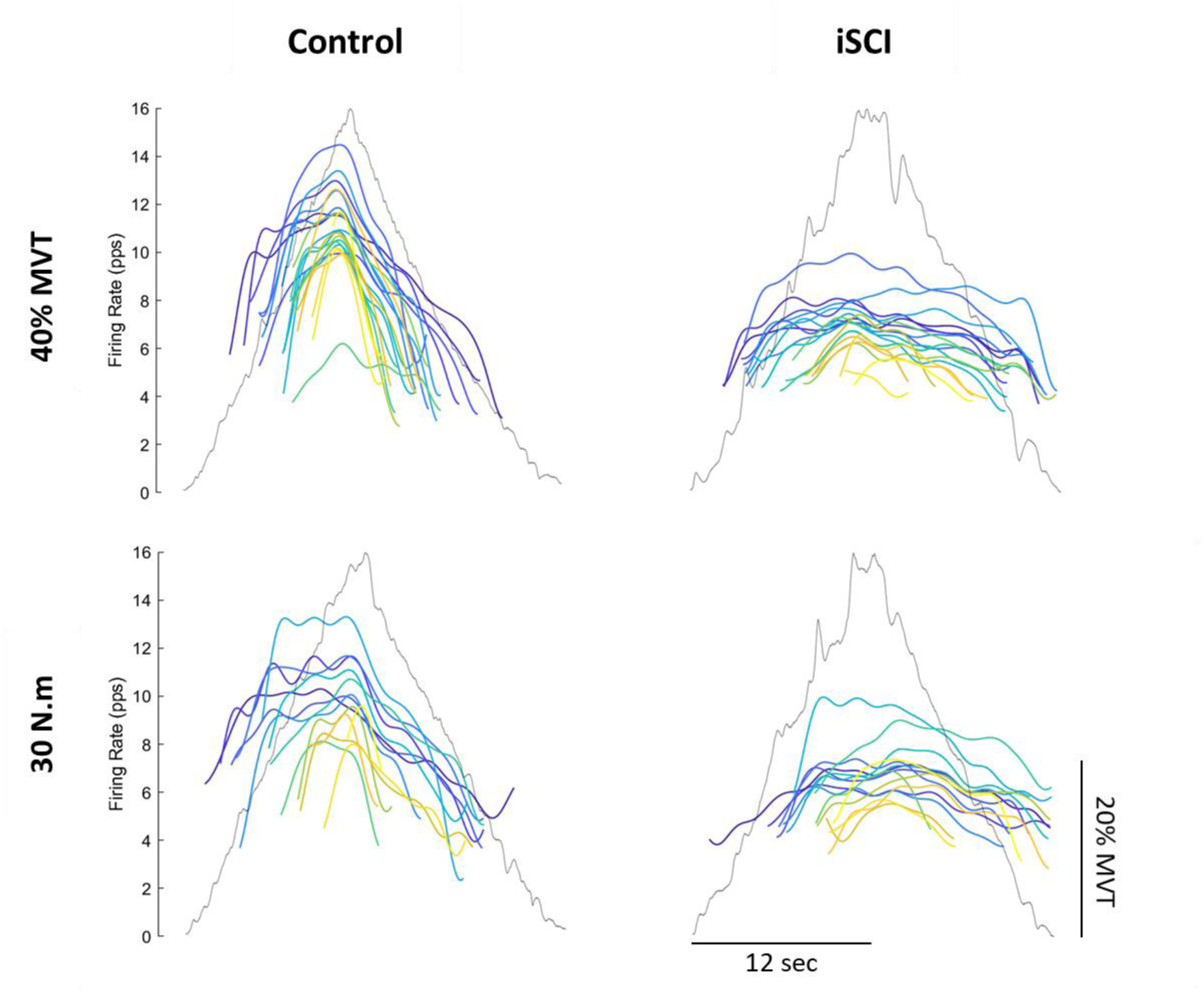
Typical motoneuron firing rates in control (CTL) and incomplete SCI (iSCI) participants. Plantarflexion torque (grey trace) and smoothed firing rates of 20 identified motoneurons (coloured traces) are shown for a control participant (left) and an individual with incomplete SCI (#10, right). Within each plot, smoothed firing rates are shown with darker colours (purple) for lower threshold neurons and lighter colours (yellow) for higher threshold neurons.

There were no significant between-group differences in recruitment threshold (p = 0.2406 for the 40% MVT contractions, and p = 0.2517 for the 30 N.m contractions), nor any interaction between group and muscle (p = 0.5848 for the 40% MVT contractions, and p = 0.2428 for the 30 N.m contractions) (Fig. 4). For the 40% MVT contractions, there was a significant interaction between group and muscle for both firing rate at recruitment (p = 0.0431) and peak firing rate (p = 0.0091). Specifically, firing rate at recruitment was lower in the incomplete SCI group than in the control group by 1.87 pps in the GM (95% CI: [1.01–2.72], d = 1.43, p = 0.0001) and by 0.88 pps in the SOL (95% CI: [0.23–1.54], d = 0.68, p = 0.0096). Peak firing rate was also lower in the incomplete SCI group by 4.52 pps in the GM (95% CI: [2.88–6.16], d = 3.27, p < 0.0001) and by 2.49 pps in the SOL (95% CI: [1.49–3.49], d = 1.8, p < 0.0001). There was no significant interaction between group and muscle for firing rate modulation (p = 0.0674) but there was a main effect of group, with lower modulation values in the incomplete SCI group (d = 1.43, p < 0.0001). Overall, similar results were observed for the 30 N.m contractions, with all firing-rate metrics lower in the incomplete SCI group compared to the control group. Specifically, there were significant main effects of group for firing rate at recruitment (d = 0.97, p = 0.0106), peak firing rate (d = 1.82, p = 0.0016), and firing rate modulation (d = 0.66, p = 0. 0147), with no interaction between group and muscle (all p > 0.4827).

**Figure 4.**
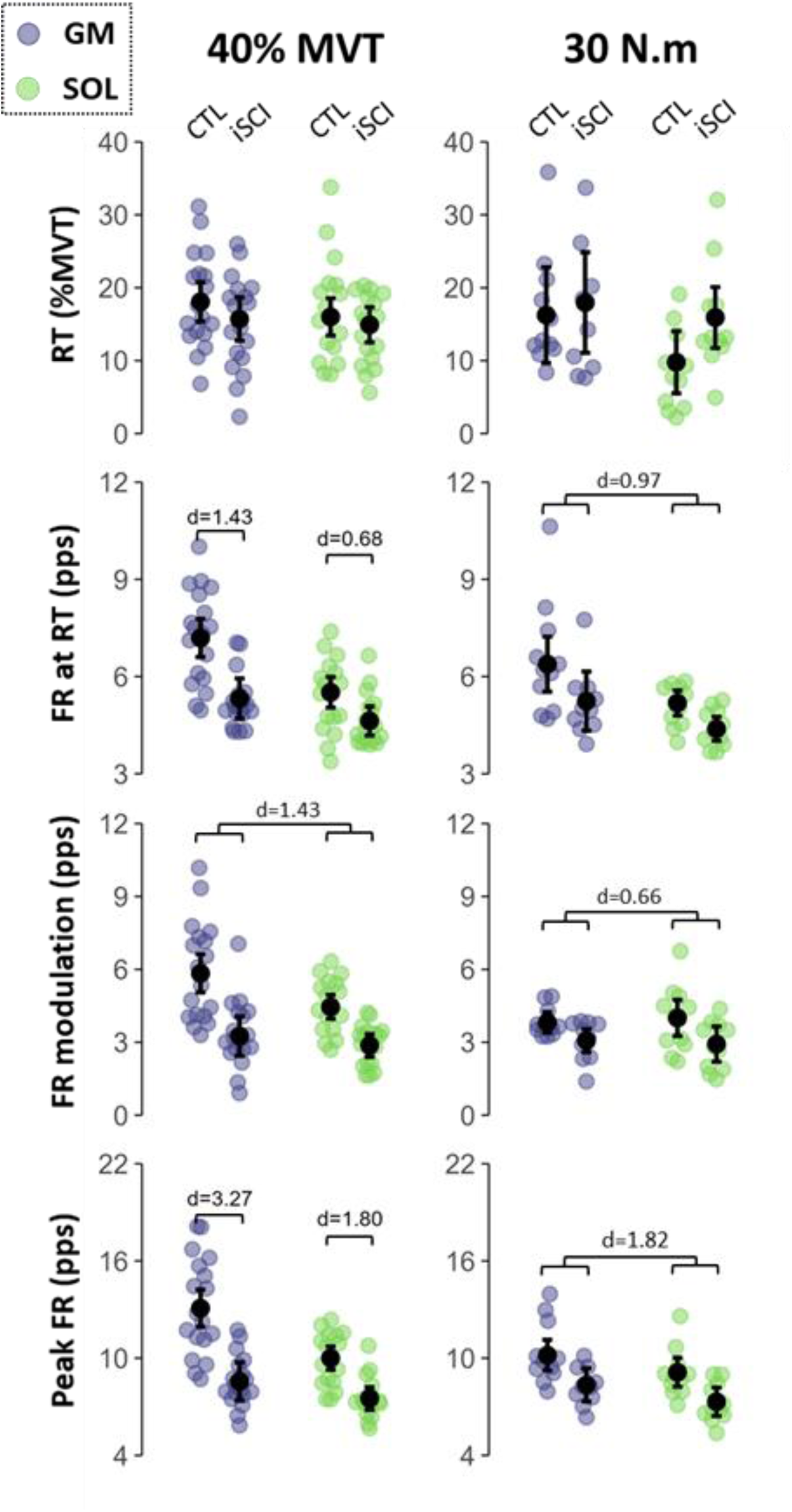
Motoneuron recruitment thresholds and rate coding. The incomplete SCI (iSCI) and control (CTL) groups are compared at either the same relative torque (40% MVT) or the same absolute torque (30 N.m). Recruitment threshold (RT), firing rate at recruitment (DR RT), firing rate modulation (FR modulation), and peak firing rate (max DR), were obtained from the gastrocnemius medialis (GM, purple) and soleus (SOL, green). Colored dots represent participant averages, while black dots and bars indicate the estimated marginal means with 95% confidence intervals predicted by linear mixed-effects models. Cohen’s d is reported for significant differences.

When considering the MUAP amplitude during the 30 N.m contractions, there was no significant between-group differences (p = 0.7115), nor any interaction between group and muscle (p = 0.0792). For the 40% MVT contractions, there was a significant interaction between group and muscle (p = 0.0424), with lower MUAP amplitude in the incomplete SCI group in the GM by 8.2 mV (95% CI: [1.05–15.38], d = 0.74, p < 0.0259) but not in the SOL (p = 0.9584).

#### Estimates of synaptic inputs to motoneurons

Brace height did not differ between groups (p = 0.3311 for the 40% MVT contractions and p = 0.2399 for the 30 N·m contractions), nor was there any interaction between group and muscle (p = 0.5974 and p = 0.3818, respectively), suggesting no group differences in estimated neuromodulatory input (Fig. 5). Conversely, attenuation slope showed no significant interaction between group and muscle (p = 0.3982), but a significant main effect of group for the 40% MVT contractions, with lower attenuation slopes in the incomplete SCI group (d = 0.91, p < 0.0001). This finding indicates a less pronounced push-pull inhibitory pattern in the incomplete SCI group compared with controls. There was no significant differences including the group factor for the 30 N.m contractions (muscle-group interaction: p = 0.2728; group effect: p = 0.2275). When considering acceleration, there was a significant interaction between group and muscle for the 40% MVT contractions (p = 0.0111). Specifically, acceleration was lower in the incomplete SCI group by 1.19 pps.s^-1^ in the GM (95% CI: [0.82–1.55], d = 1.66, p < 0.0001) and by 0.69 pps.s^-1^ in the SOL (95% CI: [0.43–0.96], d = 0.97, p < 0.0001). Similar results were observed for the 30 N.m contractions, with no muscle-group interaction (p = 0.9160) but a significant main effect of group, with lower acceleration in the incomplete SCI group (d = 0.58, p = 0.0023). Because attenuation slope differed between groups whereas brace height did not, the reduced acceleration found here likely reflects greater inhibitory input rather than altered neuromodulation.

**Figure 5.**
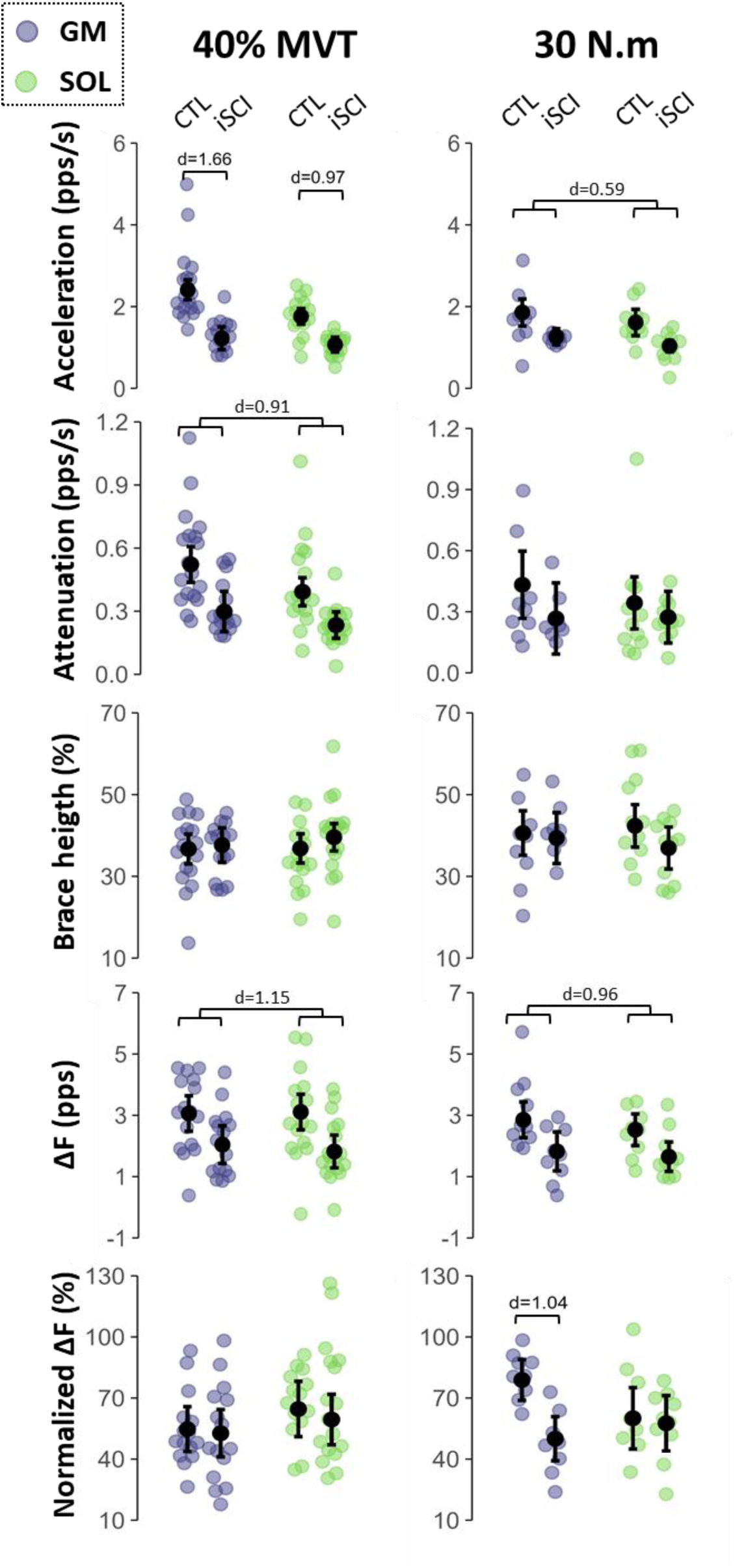
Estimates of synaptic inputs to motoneurons. The incomplete SCI (iSCI) and control (CTL) groups are compared at either the same relative torque (40% MVT) or the same absolute torque (30 N.m). The acceleration slope, attenuation slope, brace height, ΔF, and normalised ΔF were obtained from the gastrocnemius medialis (GM, purple) and soleus (SOL, green). Colored dots represent participant averages, while black dots and bars indicate the estimated marginal means with 95% confidence intervals predicted by linear mixed-effects models. Cohen’s d is reported for significant results.

When considering ΔF, there was a significant main effect of group for both the 40% MVT contractions (p = 0.0018) and the 30 N.m contractions (p = 0.0050), with no significant interaction between group and muscle (p = 0. 5350 for the 40% MVT contractions, and p = 0.7365 for the 30 N.m contractions). Specifically, ΔF was lower in the incomplete SCI group (40% MVT contraction: d = 1.2, p = 0.0017; 30 N.m contraction: p = 0.96, p = 0.0048). When considering normalised ΔF, there was no significant group effect for the 40% MVT contractions (p = 0.5504) nor significant interaction between group and muscle (p = 7940). However, for the 30 N.m contractions, a significant interaction between group and muscle was observed (p = 0.0151), with normalised ΔF being 28.8 percentage points lower in the GM of the incomplete SCI group (95% CI: [14.1–43.6], d = 1.2, p = 0.0011), while no group difference was found in the SOL (p = 0.8010).

### Modulation of motoneuron behavior in response to change in muscle length and vibration

Modulation of motoneuron firing rates was compared between the incomplete SCI and control groups for the 40% MVT contractions under two conditions: change in muscle length and the application of vibration to the tendons of the antagonist muscles. For the muscle-length condition, we compared the short and intermediate positions. For the vibration condition, we compared contractions with and without vibration at the short muscle length, as no modulation of firing rate was observed at the intermediate position with vibration in either group.

An example of smoothed motoneuron firing rate from one control and one participant with incomplete SCI (#10) at short muscle length, intermediate muscle length, and during tendon vibration is shown in Figure 6.

**Figure 6.**
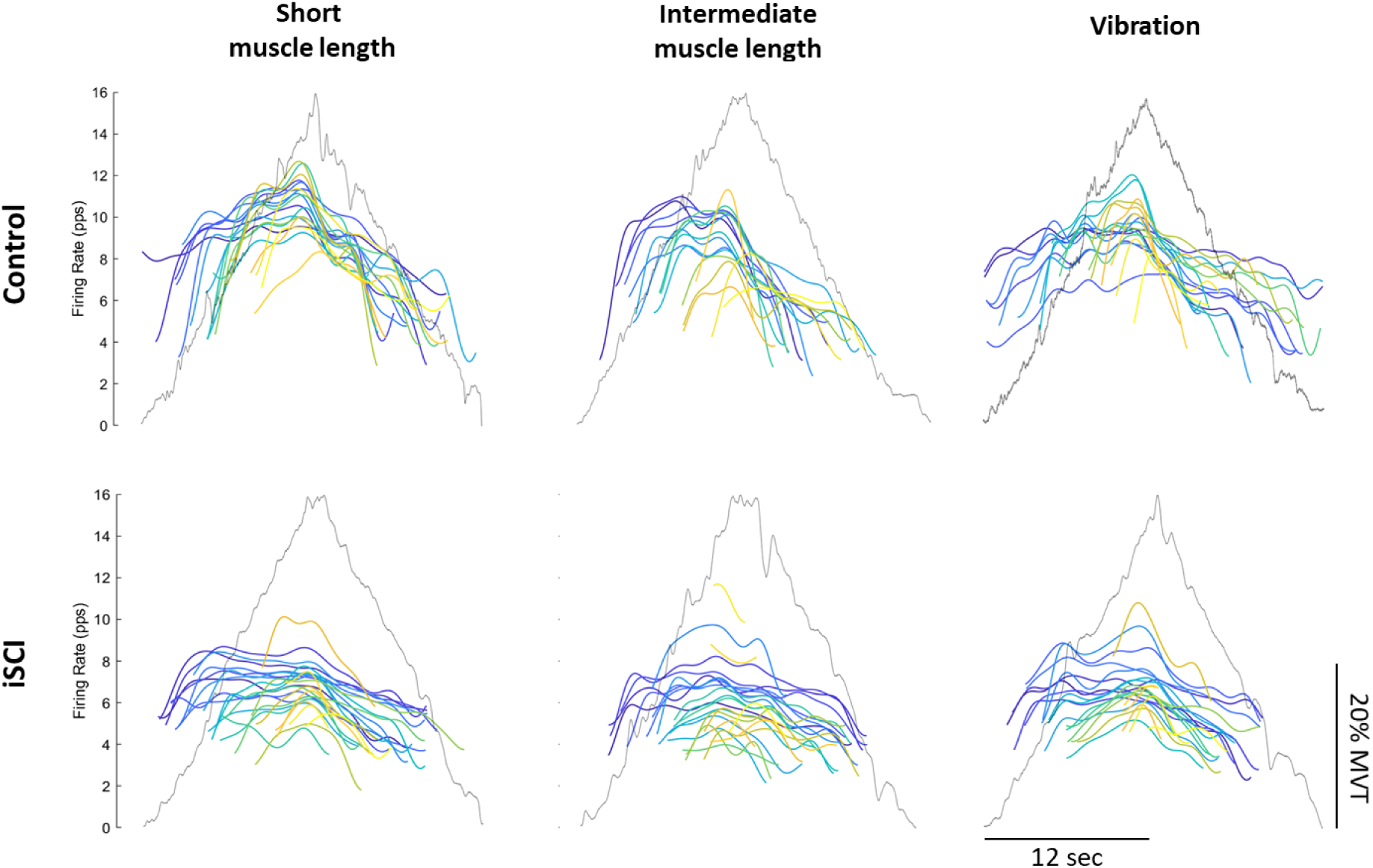
Typical effects of muscle length and vibration on motoneuron firing rates in control (CTL) and incomplete SCI (iSCI). Plantarflexion torques (grey trace) and smoothed firing rates of decomposed soleus motoneurons (colored traces) for a control participant (upper panel) and an incomplete SCI participant (#10, lower panel). Contractions were performed at short muscle length (left), intermediate muscle length (middle), and with the application of vibration to the tendons of the antagonist muscles (right). For both participants, smoothed firing rates are shown with darker colors (purple) at lower thresholds and lighter colors (yellow) at higher thresholds.

#### Recruitment threshold and rate coding

For the following results, the effects of interest are those involving interactions between group and condition (muscle length or vibration), including both group-condition interactions and group-condition-muscle interactions.

For the muscle length condition, there were no significant interactions involving group for recruitment threshold (all p > 0.0558) or firing rate at recruitment (all p > 0.0611) (Fig. 7). When considering peak firing rate, there was an interaction between group and muscle length (p = 0.0132), with no significant interaction between group, muscle length, and muscle (p = 0.1500). Specifically, the control group showed a higher peak firing rate at short compared to intermediate muscle length, regardless of the muscle (d = 0.73, p = 0.0001), whereas no difference between muscle lengths was observed in the incomplete SCI group (p = 0.5150). When considering firing rate modulation, there was a significant interaction between group, muscle length and muscle (p = 0.0144). In the control group, firing rate modulation was higher in the GM at short compared with intermediate muscle length by 0.85 pps (95% CI: [0.35–1.34], d = 0.59, p = 0.0017), whereas no difference was observed in the SOL (p = 0.7812). In the incomplete SCI group, firing rate modulation did not differ across muscle lengths, regardless of the muscle (all p > 0.3726).

**Figure 7.**
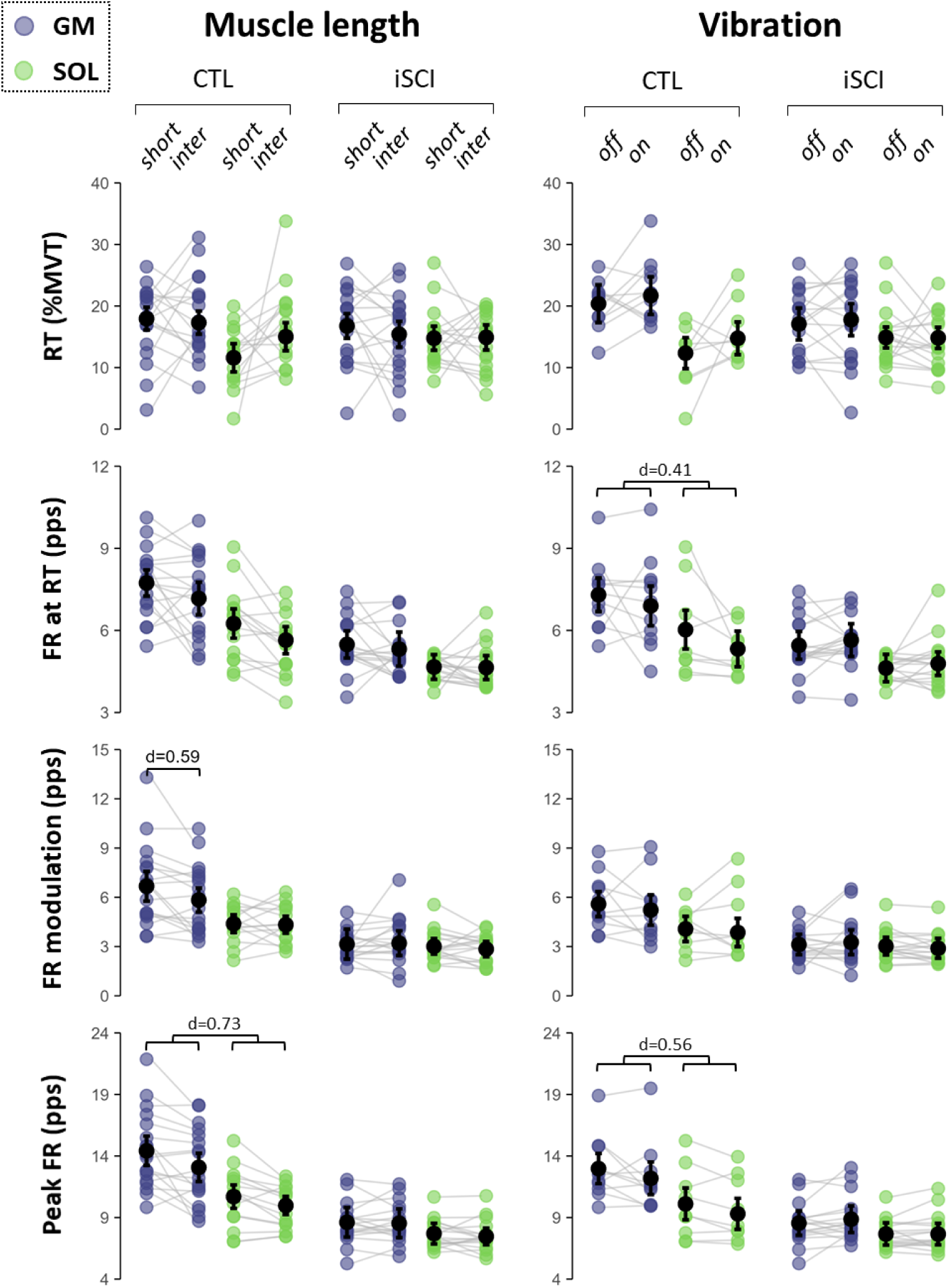
Effect of muscle length and vibration in motoneuron recruitment threshold and rate coding. The modulation of motoneuron rate coding and recruitment threshold between the incomplete SCI (iSCI) and control (CTL) groups are compared in response to changes in muscle length at the same relative torque (40% MVT), and the application of tendon vibration to antagonist muscles, at short muscle length (20° of plantarflexion). Recruitment threshold (RT), firing rate at recruitment (DR RT), firing rate modulation (FR modulation), and peak firing rate (peak FR), were obtained from the gastrocnemius medialis (GM, purple) and soleus (SOL, green). Colored dots represent participant averages, while black dots and bars indicate the estimated marginal means with 95% confidence intervals predicted by linear mixed-effects models. Cohen’s d is reported for significant results (p < 0.05).

For the vibration condition, there were no significant interactions involving the group factor for recruitment threshold (all p > 0.1719). There was an interaction between group and vibration for firing rate at recruitment (p = 0.0292) and for peak firing rate (p = 0.0029), with no significant interaction between group, vibration, and muscle (all p > 0.5268). In controls, vibration reduced firing rate at recruitment (d = 0.41, p = 0.0358) and peak firing rate (d = 0.56, p = 0.0021), whereas no significant effects were observed in the incomplete SCI group. There were no significant interactions involving the group factor for rate modulation (all p > 0.2824).

#### Estimates of synaptic inputs to motoneurons

For the muscle length condition, there was a significant interaction between group, muscle length, and muscle for brace height (p = 0.0345), although post hoc tests revealing no significant differences (all p > 0.1342). There was no interaction between group and muscle length, nor between group, muscle length, and muscle for acceleration (p > 0.1820), or attenuation (p > 0.1693) (Fig. 8). When considering ΔF, there was an interaction between group, muscle and muscle length (p = 0.0097). Specifically, there were no significant changes in the incomplete SCI group (all p > 0.2965), whereas the control group showed higher ΔF at short muscle length in the GM by 0.33 pps (95% CI: [0.019–0.64], d = 0.36, p = 0.0388) and lower ΔF at short muscle length in the SOL by 0.51 pps (95% CI: [0.026–1.0], d = 0.56, p = 0.0400). By contrast, when considering normalised ΔF, there was no significant interaction was found between group and muscle length (p = 0.8854), nor between group, muscle length, and muscle (p = 0.0938).

**Figure 8.**
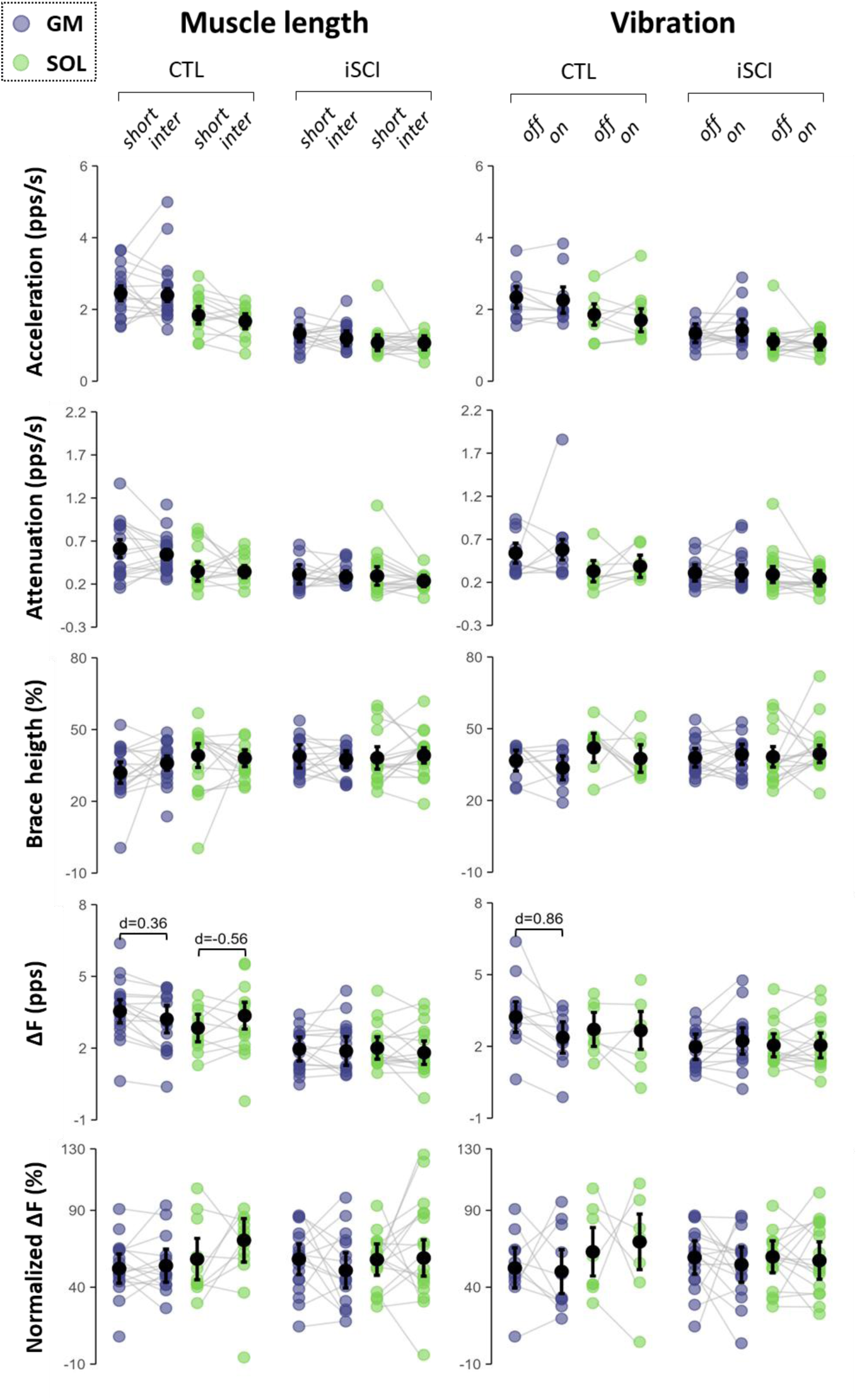
Effect of muscle length and vibration in estimates of synaptic inputs to motoneurons. The modulation of motoneuron estimates of synaptic inputs to motoneurons between the incomplete SCI (iSCI) and control (CTL) groups are compared in response to changes in muscle length at the same relative torque (40% MVT), and the application of tendon vibration to antagonist muscles, at short muscle length (20° of plantarflexion). The acceleration slope, attenuation slope, brace height, ΔF, and normalised ΔF were obtained from the gastrocnemius medialis (GM, purple) and soleus (SOL, green). Colored dots represent participant averages, while black dots and bars indicate the estimated marginal means with 95% confidence intervals predicted by linear mixed-effects models. Cohen’s d is reported for significant results (p < 0.05).

For the vibration condition, there was no interaction between group and vibration, nor between group, muscle length, and muscle for brace height (all p > 0.0584), acceleration (all p > 0.1322), or attenuation (all p > 0.2775). When considering ΔF, there was an interaction between group, muscle and vibration (p = 0.0004). Specifically, there were no significant difference in the incomplete SCI group (all p > 0.1881), whereas the control group showed higher ΔF without than with vibration in the GM by 0.79 pps (95% CI: [0.062–1.53], d = 0.86, p = 0.0005), with no difference in the SOL (p = 0.8545). By contrast, when considering normalised ΔF, there was no significant interaction was found between group and vibration (p = 0. 5309), nor between group, vibration, and muscle (p = 0.6477).

#### Results from supplementary analyses

First, because plantarflexion torque-generating capacity increases from short to intermediate muscle lengths, we included an additional contraction at an intermediate joint position (0°), matched to 40% MVT obtained at the short muscle length (20°), as presented in the supplemental materials. This condition yielded results consistent with those of the relative torque-matched contractions (see supplemental material). Second, a subset of participants with sufficient ankle range of motion (SCI: n = 9; control: n = 10) performed contractions at a long muscle length (10° dorsiflexion). In the control group, this supplemental condition strengthened the previously described muscle length–dependent effects, with the longer muscle length inducing even greater changes than the intermediate length, compared with the short length. This was accompanied by a significant reduction in ΔF in the SOL and a reduction in acceleration in GM with increasing muscle length. Importantly, no significant modulation of rate coding or synaptic input metrics was observed in the incomplete SCI group with this supplemental condition. Finally, analyses restricted to the motoneurons successfully tracked across conditions yielded converging results (see supplemental material).

## DISCUSSION

Based on non-invasive recordings of motoneurons innervating the GM and SOL muscles, this study demonstrates lower rate coding during voluntary contraction in individuals with incomplete SCI. Compared with control participants, these individuals exhibited lower firing rates at recruitment, lower firing rate modulation, and lower peak firing rates. This lower rate coding was associated with a shift in the inhibition-excitation balance toward greater inhibition, with no evidence of altered neuromodulatory input. In contrast to healthy participants, the rate coding of SCI individuals did not adapt to experimental conditions designed to modulate inhibitory input, including changes in muscle length and the application of vibration. Together, these findings suggest that motor impairment during voluntary contraction after incomplete SCI is associated with reduced rate coding caused by excessive inhibitory inputs.

### Evidence of lower rate coding after incomplete SCI

We observed markedly reduced rate coding in individuals with incomplete SCI across two lower limb muscles, as reflected by lower firing rates at recruitment, reduced firing rate modulation, and lower peak firing rates. These deficits were consistently observed at both relative (40% MVT) and absolute (30 N.m) contraction levels, making it unlikely that they are attributable to differences in maximal force-generating capacity between groups. Specifically, contractions performed at the same absolute torque corresponded to an average relative torque of ∼20% MVT in the control group but ∼48% MVT in the SCI group, indicating that individuals with SCI exhibited reduced rate coding despite operating at substantially higher relative intensities. Although these findings contrast with some previous reports showing no change in rate coding after SCI in the triceps brachii (Wiegner et al., 1993), they are consistent with prior observations of lower motoneuron firing rates in the first dorsal interosseous, biceps brachii, and tibialis anterior muscles (Debenham et al., 2024; Kizyte et al., 2025; Thomas et al., 2014). These discrepancies across studies suggest that the present findings may not generalise to all muscles after SCI. However, despite substantial heterogeneity in aetiology, motor capacity, and medication use, individuals with incomplete SCI showed consistent differences compared with the control group, supporting the generalisability of our conclusion.

### Inhibition-excitation balance contributes to the lower rate coding after incomplete SCI

To understand the mechanisms underlying the lower rate coding in individuals with SCI, we applied a recently developed geometric analysis of motoneuron firing patterns (Beauchamp et al., 2023; Chardon et al., 2024) together with the classical paired motor units analysis (Gorassini et al., 2002), to assess the contribution of synaptic inputs. As described in the Methods section, these approaches enable the estimation of the level of neuromodulation (i.e., the level of PICs in motoneurons) using brace height, as well as PIC effects to motoneuron firing. Specifically, PICs lead to an initial amplification effect on firing, quantified by the acceleration slope, and a prolongation effect, quantified by ΔF (see Methods). Importantly, brace height reflects neuromodulation because it is independent of the level of inhibition, unlike the two other metrics. This distinction reflects the ability of inhibitory inputs to dampen the effects of neuromodulation on firing. Our result that brace height did not differ between groups suggest preserved neuromodulatory inputs in individuals with SCI. In contrast, both the acceleration slope and ΔF were consistently lower in the incomplete SCI group, irrespective of contraction intensity or muscle. Taken together, these findings suggest that PIC amplification and prolongation effects on motoneuron firing are reduced following incomplete SCI, despite preserved neuromodulatory input. Importantly, comparisons of PIC effects across motoneuron populations with different modulation of firing rate are inherently challenging, as ΔF depends on firing rate modulation of control motoneurons, which was lower in the incomplete SCI group. To address this limitation, ΔF values were normalised to the maximal ΔF values achievable relative to control unit modulation (Škarabot et al., 2025). After normalisation, group differences were no longer significantly different, except for the GM in the absolute comparison condition. A possible interpretation is that although PIC effects are lower in absolute terms after incomplete SCI, their relative contribution to motoneuron firing is preserved, supporting PICs as a continuing source of depolarising current after incomplete SCI. Of note, the normalisation procedure applied to ΔF likely makes it a more neuromodulation-specific indicator by removing the influence of inhibition on firing, similar to brace height (Beauchamp et al., 2023; Oya et al., 2009).

The geometric analysis of motoneuron firing patterns also allows to estimate the balance between inhibitory and excitatory inputs (referred to as the pattern of inhibition) through the attenuation slope (Beauchamp et al., 2023; Chardon et al., 2024). Attenuation slopes were lower in the incomplete SCI group for both GM and SOL at 40% MVT, suggesting an altered inhibitory pattern, characterised by a greater inhibitory influence. The pattern of inhibition can range from a strong positive covariation between excitation and inhibition (proportional pattern) to a strong negative covariation (push-pull pattern). The push-pull pattern has been proposed as a mechanism by which the central nervous system increases the input-output gain of motoneurons by raising background inhibition followed by disinhibition to increase force production (Johnson et al., 2012). This pattern is supported by spinal circuits, such as the organisation of Ia afferents and Ia inhibitory interneurons (Johnson et al., 2012), and aligns with firing patterns observed in healthy humans (Chardon et al., 2024). A push-pull pattern is required to attain high firing rates during high-force contractions (Škarabot et al., 2025) and is more pronounced in trained than in untrained individuals (Škarabot et al., 2026). The lower attenuation slope observed in the incomplete SCI group is consistent with the behavior predicted by simulations assuming a more proportional inhibitory-excitatory pattern (Chardon et al., 2024). Accordingly, a reduced capacity to regulate inhibition during increasing force production likely contributes to altered motoneuron rate coding. Furthermore, the absence of firing rate modulation in response to experimental manipulations of inhibitory input (Fig. 7), achieved through changes in muscle length or the application of vibration, suggests that individuals with incomplete SCI may already operate under elevated inhibition during voluntary contractions, consistent with a more proportional inhibitory-excitatory pattern. One possible mechanism underlying this elevated inhibition is impaired regulation of recurrent inhibition, resulting in increased inhibition concomitant with increasing excitation. In healthy individuals, descending control modulates the activity of Renshaw cells during voluntary contraction. However, this control is often disrupted after upper motor neuron lesions (Katz & Pierrot-Deseilligny, 1982), as a result of decreased supraspinal inhibitory control (Jankowska, 1992). This interpretation is further supported by previous evidence of enhanced recurrent inhibition following incomplete SCI (Shefner et al., 1992).

### Lack of motoneuron behavior adaptation to experimental manipulation of inhibitory inputs

We examined the adaptation of motoneurons by experimentally manipulating inhibitory inputs, through changes in muscle length via joint position and by applying vibration to the tendons of the antagonist muscles during submaximal voluntary contractions. Specifically, increased muscle length was used to notably elicit recurrent inhibition (Colard et al., 2025), whereas antagonist tendon vibration was used to generate reciprocal inhibition through Ia inhibitory interneurons (Burke et al., 1976). As expected, the control group exhibited reduced firing rates at longer muscle lengths and during tendon vibration, accompanied by a reduction in the PIC prolongation effect, especially in the GM. These results largely align with previous work on the effects of muscle length (Beauchamp et al., 2025; Goreau et al., 2025; Valenčič et al., 2026) and antagonist tendon vibration (Mesquita et al., 2022; Pearcey et al., 2022) and are likely explained by the dampening effect of synaptic inhibition on motoneuron PICs (Bui et al., 2008; Kuo et al., 2003)

In contrast to the control group, the incomplete SCI group showed no change in rate coding in response to changes in muscle length or vibration (Fig. 7). One possible explanation for this lack of modulation is that the tested paradigms did not further increase the level of inhibitory inputs in SCI participants, likely because inhibitory interneurons operate near their maximal activity even at very low contraction intensities during voluntary contraction. This is consistent with the high level of inhibition associated with the more proportional inhibitory–excitatory pattern, as described above. Alternatively, the experimental paradigms employed may not have sufficiently activated inhibitory interneurons to further enhance inhibitory inputs. Indeed, reciprocal inhibition is replaced by facilitation after SCI (Crone, 2003; Knikou & Mummidisetty, 2011), and the modulation of recurrent inhibition with muscle length is likely dependent on descending control (Colard et al., 2025), which is altered after SCI (Katz & Pierrot-Deseilligny, 1982; Shefner et al., 1992). Other potential mechanisms that could have limited an increase in inhibitory inputs include the loss of monoaminergic inputs that regulate Ia inhibitory interneurons (Hammar & Jankowska, 2003; Jankowska et al., 2000), as well as a downregulation of KCC2, which increases inhibitory input strength and is reduced after SCI (Boulenguez et al., 2010). Regardless of the underlying mechanisms, these findings highlight the limited capacity of individuals with SCI to adapt the motor command via inhibitory inputs.

Of note, our findings of increased inhibition during voluntary contraction stand in clear contrast to the mechanisms underlying muscle spasms at rest after SCI, which are characterised by reduced synaptic inhibition and increased PICs (Gorassini et al., 2004; Mahrous et al., 2024). An impaired capacity to regulate inhibition according to task demands (e.g., maintaining muscle relaxation or increasing muscle contraction) may explain this discrepancy and is consistent with our findings of a limited capacity in individuals with SCI to adapt the motor command via inhibitory inputs.

### Reduced rate coding is compensated by the recruitment of a greater number of motor units after incomplete SCI

Despite exhibiting lower rate coding than control participants, individuals with incomplete SCI were able to produce the same absolute torque (30 N.m). The lower firing rate observed in the incomplete SCI group must therefore have been compensated by the recruitment of larger motor units and/or a greater number of motor units. Due to limitations of EMG decomposition in reliably assessing recruitment, we cannot directly determine the compensatory strategies employed. Assuming similar spatial sampling of motor units and comparable volume conductor effects across populations, we used MUAP amplitude as a proxy for motor unit size. We found no between-group differences at 30 N.m, suggesting that individuals with incomplete SCI compensated for their lower rate coding primarily through the recruitment of additional motor units, which is further supported the observation of higher EMG amplitudes in the SCI group (Fig. 1C). This interpretation is consistent with studies showing that motoneurons are recruited up to higher force levels in individuals with SCI than in controls (Thomas et al., 1997; Zijdewind & Thomas, 2003). Importantly, this interpretation is based on indirect measures and unverifiable assumptions; and further work is needed to confirm a greater reliance on recruitment in individuals with incomplete SCI.

### Implications for motor impairments related to incomplete SCI

Our results showing that individuals with incomplete SCI operate under elevated inhibitory inputs during voluntary contraction may help explain paresis. Indeed, compensating for this inhibition would require a disproportionate increase in excitatory input to further raise motoneuron output. Given that excitatory descending inputs are already compromised after SCI, this compensatory capacity is likely limited. Consequently, excitatory drive may reach saturation at relatively low force levels, thereby constraining maximal torque production through reduced voluntary activation (Kim et al., 2015).

In addition, unlike the control group, the incomplete SCI group showed no change in rate coding in response to changes in muscle length. This adaptive pattern in healthy individuals may represent a mechanism that adjusts neural drive to the diminished muscle force-generating capacity at shorter lengths (Goreau et al., 2025). Conversely, the failure to adapt rate coding after incomplete SCI may lead to muscle length-dependent paresis. While this phenomenon has not been directly quantified after SCI, it is consistent with the concept of spastic paresis syndrome, common to central nervous system disorders (Baude et al., 2019; Gracies, 2005), and observed, for example, after stroke (Vinti et al., 2015). In line with this, incomplete SCI is associated with reduced central activation (i.e., paresis), which is more pronounced during shortening contractions (Kim et al., 2015).

Finally, compensation for reduced rate coding through the recruitment of a greater number of motor units after incomplete SCI implies increased use of high-threshold motor units, including fast fatigable units. This likely contributes to the increased fatigability observed after incomplete SCI (Thomas & Del Valle, 2001).

## Conclusion

This study provides the first evidence in humans that voluntary motor deficits after incomplete SCI may be associated with a disrupted balance between inhibitory and excitatory synaptic inputs to motoneurons. During voluntary contractions, individuals with SCI show a limited capacity to modulate inhibitory inputs, resulting in reduced spinal motoneuron rate coding. These findings support excessive inhibition as a key neural mechanism contributing to muscle weakness and impaired motor function after incomplete SCI. Clarifying the mechanisms responsible for this excessive inhibition represents a critical next step, as it constitutes a promising therapeutic target for restoring force-generating capacity after incomplete SCI.

## DATA AVAILABILITY

Dataset of analysed data is available at 10.6084/m9.figshare.31338631

## FOUNDINGS

This study was partly funded by a grant from the French National Research Agency (ANR-24-CE17-5805; Neuromotor project). François Hug is supported by the French government, through the UCAJEDI Investments in the Future project managed by the ANR with the reference number ANR-15-IDEX-01, by a fellowship from the Institut Universitaire de France (IUF).

## REFERENCES

Avrillon, S., Hug, F., Baker, S. N., Gibbs, C., & Farina, D. (2024). Tutorial on MUedit : An open-source software for identifying and analysing the discharge timing of motor units from electromyographic signals. Journal of Electromyography and Kinesiology, 77, 102886. 10.1016/j.jelekin.2024.102886

Avrillon, S., Hug, F., Enoka, R. M., Caillet, A. H., & Farina, D. (2024). The identification of extensive samples of motor units in human muscles reveals diverse effects of neuromodulatory inputs on the rate coding. eLife, 13, RP97085. 10.7554/eLife.97085.3

Baude, M., Nielsen, J. B., & Gracies, J.-M. (2019). The neurophysiology of deforming spastic paresis : A revised taxonomy. Annals of Physical and Rehabilitation Medicine, 62(6), 426-430. 10.1016/j.rehab.2018.10.004

Beauchamp, J. A., Khurram, O. U., Dewald, J. P. A., Heckman, C., & Pearcey, G. E. P. (2022). A computational approach for generating continuous estimates of motor unit discharge rates and visualizing population discharge characteristics. Journal of Neural Engineering, 19(1), 016007. 10.1088/1741-2552/ac4594

Beauchamp, J. A., Pearcey, G. E. P., Khurram, O. U., Chardon, M., Wang, Y. C., Powers, R. K., Dewald, J. P. A., & Heckman, C. (2023). A geometric approach to quantifying the neuromodulatory effects of persistent inward currents on individual motor unit discharge patterns. Journal of Neural Engineering, 20(1), 016034. 10.1088/1741-2552/acb1d7

Beauchamp, J. A., Pearcey, G. E. P., Khurram, O. U., Negro, F., Dewald, J. P. A., & Heckman, C. J. (2025). Intrinsic properties of spinal motoneurons degrade ankle torque control in humans. The Journal of Physiology, 603(8), 2443-2463. 10.1113/JP287446

Bennett, D. J., Hultborn, H., Fedirchuk, B., & Gorassini, M. (1998). Short-Term Plasticity in Hindlimb Motoneurons of Decerebrate Cats. Journal of Neurophysiology, 80(4), 2038-2045. 10.1152/jn.1998.80.4.2038

Boulenguez, P., Liabeuf, S., Bos, R., Bras, H., Jean-Xavier, C., Brocard, C., Stil, A., Darbon, P., Cattaert, D., Delpire, E., Marsala, M., & Vinay, L. (2010). Down-regulation of the potassium-chloride cotransporter KCC2 contributes to spasticity after spinal cord injury. Nature Medicine, 16(3), 302-307. 10.1038/nm.2107

Bui, T. V., Grande, G., & Rose, P. K. (2008). Relative Location of Inhibitory Synapses and Persistent Inward Currents Determines the Magnitude and Mode of Synaptic Amplification in Motoneurons. Journal of Neurophysiology, 99(2), 583-594. 10.1152/jn.00718.2007

Burke, D., Hagbarth, K. E., Löfstedt, L., & Wallin, B. G. (1976). The responses of human muscle spindle endings to vibration of non-contracting muscles. The Journal of Physiology, 261(3), 673-693. 10.1113/jphysiol.1976.sp011580

Chardon, M. K., Wang, Y. C., Garcia, M., Besler, E., Beauchamp, J. A., D’Mello, M., Powers, R. K., & Heckman, C. J. (2024). Supercomputer framework for reverse engineering firing patterns of neuron populations to identify their synaptic inputs. eLife, 12, RP90624. 10.7554/eLife.90624

Colard, J., Duclay, J., Betus, Y., Cattagni, T., & Jubeau, M. (2025). Muscle Length Modulates Recurrent Inhibition and Presynaptic Inhibition of Ia Afferents Differently Depending on Type of Contraction. European Journal of Neuroscience, 62(1), e70172. 10.1111/ejn.70172

Crone, C. (2003). Appearance of reciprocal facilitation of ankle extensors from ankle flexors in patients with stroke or spinal cord injury. Brain, 126(2), 495-507. 10.1093/brain/awg036

D’Amico, J. M., Condliffe, E. G., Martins, K. J. B., Bennett, D. J., & Gorassini, M. A. (2014). Recovery of neuronal and network excitability after spinal cord injury and implications for spasticity. Frontiers in Integrative Neuroscience, 8. 10.3389/fnint.2014.00036

Debenham, M. I. B., Franz, C. K., & Berger, M. J. (2024). Neuromuscular consequences of spinal cord injury : New mechanistic insights and clinical considerations. Muscle & Nerve, 70(1), 12-27. 10.1002/mus.28070

Del Vecchio, A., Holobar, A., Falla, D., Felici, F., Enoka, R. M., & Farina, D. (2020). Tutorial : Analysis of motor unit discharge characteristics from high-density surface EMG signals. Journal of Electromyography and Kinesiology, 53, 102426. 10.1016/j.jelekin.2020.102426

Gorassini, M. A. (2004). Role of motoneurons in the generation of muscle spasms after spinal cord injury. Brain, 127(10), 2247-2258. 10.1093/brain/awh243

Gorassini, M., Yang, J. F., Siu, M., & Bennett, D. J. (2002). Intrinsic Activation of Human Motoneurons : Possible Contribution to Motor Unit Excitation. Journal of Neurophysiology, 87(4), 1850-1858. 10.1152/jn.00024.2001

Goreau, V., Morvan, Q., Hug, F., Le Sant, G., Gross, R., & Cattagni, T. (2025). Modulation of persistent inward currents in alpha motoneurons with joint angle depends on muscle length. Journal of Neurophysiology, 134(2), 493-503. 10.1152/jn.00097.2025

Gracies, J.-M. (2005). Pathophysiology of spastic paresis. II : Emergence of muscle overactivity. Muscle & Nerve, 31(5), 552-571. 10.1002/mus.20285

Hammar, I., & Jankowska, E. (2003). Modulatory Effects of α1-, α2-, and β-Receptor Agonists on Feline Spinal Interneurons with Monosynaptic Input from Group I Muscle Afferents. The Journal of Neuroscience, 23(1), 332-338. 10.1523/JNEUROSCI.23-01-00332.2003

Hassan, A., Thompson, C. K., Negro, F., Cummings, M., Powers, R. K., Heckman, C. J., Dewald, J. P. A., & McPherson, L. M. (2020). Impact of parameter selection on estimates of motoneuron excitability using paired motor unit analysis. Journal of Neural Engineering, 17(1), 016063. 10.1088/1741-2552/ab5eda

Hug, F., Avrillon, S., Del Vecchio, A., Casolo, A., Ibanez, J., Nuccio, S., Rossato, J., Holobar, A., & Farina, D. (2021). Analysis of motor unit spike trains estimated from high-density surface electromyography is highly reliable across operators. Journal of Electromyography and Kinesiology: Official Journal of the International Society of Electrophysiological Kinesiology, 58, 102548. 10.1016/j.jelekin.2021.102548

Hultborn, H., Denton, M. E., Wienecke, J., & Nielsen, J. B. (2003). Variable amplification of synaptic input to cat spinal motoneurones by dendritic persistent inward current. The Journal of Physiology, 552(3), 945-952. 10.1113/jphysiol.2003.050971

Hyngstrom, A. S., Johnson, M. D., Miller, J. F., & Heckman, C. J. (2007). Intrinsic electrical properties of spinal motoneurons vary with joint angle. Nature Neuroscience, 10(3), 363-369. 10.1038/nn1852

Jankowska, E. (1992). Interneuronal relay in spinal pathways from proprioceptors. Progress in Neurobiology, 38(4), 335-378. 10.1016/0301-0082(92)90024-9

Jankowska, E., Hammar, I., Chojnicka, B., & Hedén, C. H. (2000). Effects of monoamines on interneurons in four spinal reflex pathways from group I and/or group II muscle afferents. European Journal of Neuroscience, 12(2), 701-714. 10.1046/j.1460-9568.2000.00955.x

Jayaraman, A., Gregory, C. M., Bowden, M., Stevens, J. E., Shah, P., Behrman, A. L., & Vandenborne, K. (2006). Lower extremity skeletal muscle function in persons with incomplete spinal cord injury. Spinal Cord, 44(11), 680-687. 10.1038/sj.sc.3101892

Johnson, M. D., Hyngstrom, A. S., Manuel, M., & Heckman, C. J. (2012). Push–Pull Control of Motor Output. The Journal of Neuroscience, 32(13), 4592-4599. 10.1523/JNEUROSCI.4709-11.2012

Katz, R., & Pierrot-Deseilligny, E. (1982). RECURRENT INHIBITION OF α—MOTONEURONS IN PATIENTS WITH UPPER MOTOR NEURON LESIONS. Brain, 105(1), 103-124. 10.1093/brain/105.1.103

Kim, H. E., Corcos, D. M., & Hornby, T. G. (2015). Increased spinal reflex excitability is associated with enhanced central activation during voluntary lengthening contractions in human spinal cord injury. Journal of Neurophysiology, 114(1), 427-439. 10.1152/jn.01074.2014

Kizyte, A., Zhang, H., Forslund, E. B., Gutierrez-Farewik, E. M., & Wang, R. (2025). Neuromuscular adaptations in soleus and tibialis anterior muscles in persons with spinal cord injury. Journal of NeuroEngineering and Rehabilitation, 22(1), 239. 10.1186/s12984-025-01794-7

Knikou, M., & Mummidisetty, C. K. (2011). Reduced reciprocal inhibition during assisted stepping in human spinal cord injury. Experimental Neurology, 231(1), 104-112. 10.1016/j.expneurol.2011.05.021

Kuo, J. J., Lee, R. H., Johnson, M. D., Heckman, H. M., & Heckman, C. J. (2003). Active Dendritic Integration of Inhibitory Synaptic Inputs In Vivo. Journal of Neurophysiology, 90(6), 3617-3624. 10.1152/jn.00521.2003

Kuznetsova, A., Brockhoff, P. B., & Christensen, R. H. B. (2017). lmerTest Package : Tests in Linear Mixed Effects Models. Journal of Statistical Software, 82(13). 10.18637/jss.v082.i13

Kuzyk, S. L., Smart, R. R., Simpson, C. L., Fedorov, A., & Jakobi, J. M. (2018). Influence of fascicle length on twitch potentiation of the medial gastrocnemius across three ankle angles. European Journal of Applied Physiology, 118(6), 1199-1207. 10.1007/s00421-018-3849-4

Li, Y., & Bennett, D. J. (2003). Persistent Sodium and Calcium Currents Cause Plateau Potentials in Motoneurons of Chronic Spinal Rats. Journal of Neurophysiology, 90(2), 857-869. 10.1152/jn.00236.2003

Mahrous, A., Birch, D., Heckman, C. J., & Tysseling, V. (2024). Muscle Spasms after Spinal Cord Injury Stem from Changes in Motoneuron Excitability and Synaptic Inhibition, Not Synaptic Excitation. The Journal of Neuroscience, 44(1), e1695232023. 10.1523/JNEUROSCI.1695-23.2023

McDonald, J. W., & Sadowsky, C. (2002). Spinal-cord injury. The Lancet, 359(9304), 417-425. 10.1016/S0140-6736(02)07603-1

Mesquita, R. N. O., Taylor, J. L., Trajano, G. S., Škarabot, J., Holobar, A., Gonçalves, B. A. M., & Blazevich, A. J. (2022). Effects of reciprocal inhibition and whole-body relaxation on persistent inward currents estimated by two different methods. The Journal of Physiology, 600(11), 2765-2787. 10.1113/JP282765

Murray, K. C., Nakae, A., Stephens, M. J., Rank, M., D’Amico, J., Harvey, P. J., Li, X., Harris, R. L. W., Ballou, E. W., Anelli, R., Heckman, C. J., Mashimo, T., Vavrek, R., Sanelli, L., Gorassini, M. A., Bennett, D. J., & Fouad, K. (2010). Recovery of motoneuron and locomotor function after spinal cord injury depends on constitutive activity in 5-HT2C receptors. Nature Medicine, 16(6), 694-700. 10.1038/nm.2160

Negro, F., Muceli, S., Castronovo, A. M., Holobar, A., & Farina, D. (2016). Multi-channel intramuscular and surface EMG decomposition by convolutive blind source separation. Journal of Neural Engineering, 13(2), 026027. 10.1088/1741-2560/13/2/026027

Oya, T., Riek, S., & Cresswell, A. G. (2009). Recruitment and rate coding organisation for soleus motor units across entire range of voluntary isometric plantar flexions : Recruitment and rate coding strategies for soleus motor units. The Journal of Physiology, 587(19), 4737-4748. 10.1113/jphysiol.2009.175695

Pearcey, G. E. P., Khurram, O. U., Beauchamp, J. A., Negro, F., & Heckman, C. J. (2022). Antagonist tendon vibration dampens estimates of persistent inward currents in motor units of the human lower limb [Preprint]. Physiology. 10.1101/2022.08.02.502526

Powers, R. K., & Heckman, C. J. (2017). Synaptic control of the shape of the motoneuron pool input-output function. Journal of Neurophysiology, 117(3), 1171-1184. 10.1152/jn.00850.2016

Revill, A. L., & Fuglevand, A. J. (2017). Inhibition linearizes firing rate responses in human motor units : Implications for the role of persistent inward currents: Inhibition linearizes firing rate responses in motor units. The Journal of Physiology, 595(1), 179-191. 10.1113/JP272823

Schwindt, P. C., & Crill, W. E. (1980). Properties of a persistent inward current in normal and TEA-injected motoneurons. Journal of Neurophysiology, 43(6), 1700-1724. 10.1152/jn.1980.43.6.1700

Shefner, J. M., Berman, S. A., Sarkarati, M., & Young, R. R. (1992). Recurrent inhibition is increased in patients with spinal cord injury. Neurology, 42(11), 2162-2162. 10.1212/WNL.42.11.2162

Škarabot, J., Beauchamp, J. A., & Pearcey, G. E. P. (2025). Human motor unit discharge patterns reveal differences in neuromodulatory and inhibitory drive to motoneurons across contraction levels. Journal of Neurophysiology, 134(5), 1429-1444. 10.1152/jn.00249.2025

Škarabot, J., Thomason, H., Nazaroff, B. M., Connelly, C. D., Valenčič, T., Ho, M. L., Tyagi, K., Beauchamp, J. A., & Pearcey, G. E. P. (2026). The modulation of human motoneuron discharge patterns with contraction force in resistance- and endurance-trained individuals. Journal of Applied Physiology, japplphysiol.00835.2025. 10.1152/japplphysiol.00835.2025

Thomas, C. K., Bakels, R., Klein, C. S., & Zijdewind, I. (2014). Human spinal cord injury : Motor unit properties and behaviour. Acta Physiologica, 210(1), 5-19. 10.1111/apha.12153

Thomas, C. K., Broton, J. G., & Calancie, B. (1997). Motor unit forces and recruitment patterns after cervical spinal cord injury. Muscle & Nerve, 20(2), 212-220. 10.1002/(SICI)1097-4598(199702)20:2%253C212::AID-MUS12%253E3.0.CO;2-4

Thomas, C. K., & Del Valle, A. (2001). The role of motor unit rate modulation versus recruitment in repeated submaximal voluntary contractions performed by control and spinal cord injured subjects. Journal of Electromyography and Kinesiology, 11(3), 217-229. 10.1016/S1050-6411(00)00055-9

Valenčič, T., Maeo, S., Kluzek, S., Holobar, A., Škarabot, J., & Folland, J. P. (2026). Motor unit discharge properties of the vastii muscles and their modulation with contraction level depend on the knee-joint angle. Journal of Applied Physiology, 140(1), 322-337. 10.1152/japplphysiol.00951.2024

Vinti, M., Bayle, N., Hutin, E., Burke, D., & Gracies, J.-M. (2015). Stretch-sensitive paresis and effort perception in hemiparesis. Journal of Neural Transmission, 122(8), 1089-1097. 10.1007/s00702-015-1379-3

Wiegner, A. W., Wierzbicka, M. M., Davies, L., & Young, R. R. (1993). Discharge properties of single motor units in patients with spinal cord injuries. Muscle & Nerve, 16(6), 661-671. 10.1002/mus.880160613

Zijdewind, I., & Thomas, C. K. (2003). Motor Unit Firing During and After Voluntary Contractions of Human Thenar Muscles Weakened by Spinal Cord Injury. Journal of Neurophysiology, 89(4), 2065-2071. 10.1152/jn.00492.2002

