## Supplemental material for "Disrupted Inhibitory-Excitatory Balance Underlies Spinal Motoneuron Dysfunction in Incomplete Spinal ord Injury"

### SUPPLEMENTARY ANALYSIS 1 : COMPARING MUSCLE LENGTHS AT SIMILAR ABSOLUTE CONTRACTION TORQUE

#### Supplementary methods 1

In the core manuscript, to assess how motoneuron behavior adapts to muscle length, participants performed contractions at an intermediate muscle length (ankle at 0°) and at a short muscle length (20° plantarflexion). These triangular contractions reached 40% of the MVT at the respective joint position. However, because plantarflexion torque-generating capacity increases from short to intermediate muscle lengths, an additional contraction was performed at the intermediate position (0°), with torque matched to 40% MVT obtained at the short muscle length (20°).

In addition, participants who were able to perform a contraction at 10° of dorsiflexion completed this condition at a torque matched to 40% MVT obtained at the short muscle length. This allowed comparison across a wider range of muscle lengths by including a long muscle length condition.

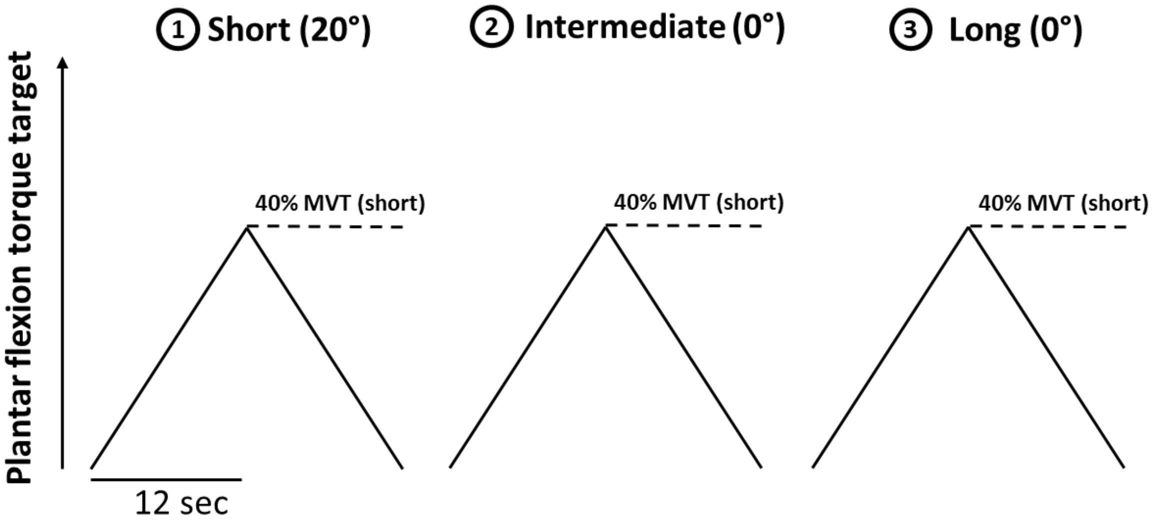


**Supplementary Figure 1. Experimental setup.** Participants performed isometric submaximal plantarflexions at three ankle positions, corresponding to different plantar flexor lengths: 20° of plantar flexion(short muscle length position), 0° (intermediate muscle length position), and 10° of dorsiflexion. The intensity of the plantarflexions followed either the same absolute torque, which correspond to 40% of the maximal voluntary contraction (MVT) in the short muscle length position.

**Supplementary results 1**

*Recruitment threshold and rate coding*

For the following results, the effects of interest are those involving interactions between group and muscle length (short, intermediate or long), including both group-muscle length interactions and group-muscle length-muscle interactions.

Participants did not perform maximal voluntary contractions at the long muscle length position, precluding determination of recruitment thresholds expressed as a percentage of MVT. Accordingly, in this condition, recruitment threshold was defined as the delay between contraction onset and the first firing. There was a significant interaction between group, muscle length, and muscle for recruitment threshold ($p > 0.0001$). Specifically, the control group showed a higher recruitment threshold in the GM at short compared with long muscle length by 1.9 s (95% CI: [0.9–3.0], $d = 0.67$, $p = 0.0002$). The incomplete SCI group showed higher recruitment threshold in the SOL at short than at long muscle length by 1.2 s (95% CI: [0.1–2.3], $d = 0.41$, $p = 0.0346$).

When considering firing rate at recruitment, there was a significant interaction between group and muscle length ($p = 0.0030$), with no significant interaction between group, muscle length, and muscle ($p = 0.9905$). Specifically, the control group showed a higher firing rate at recruitment at short compared with intermediate ($d = 0.72$, $p < 0.0001$) and long muscle length ($d = 0.90$, $p = 0.0001$), regardless of the muscle. Conversely, no difference between muscle lengths was observed in the incomplete SCI group (all $p > 0.1612$).

When considering firing rate modulation, there was a significant interaction between group and muscle length ($p = 0.0144$), with no significant interaction between group, muscle length, and muscle ($p = 0.0691$). Specifically, the control group showed a higher firing rate modulation at short compared to intermediate ($d = 0.51$, $p = 0.0019$) and long muscle length ($d = 0.92$, $p < 0.0001$), as well as a higher firing rate modulation at intermediate compared to long muscle length ($d = 0.41$, $p = 0.0203$), regardless of the muscle. Conversely, no difference between muscle lengths was observed in the incomplete SCI group (all $p > 0.1534$).

When considering peak firing rate, there was a significant interaction between group, muscle length, and muscle ($p = 0.0439$). Specifically, the control group showed a higher peak firing rate in the GM at short than at intermediate by 2.2 pps (95% CI: [1.3–3.0], $d = 1.5$, $p < 0.0001$) and long muscle length by 2.8 pps (95% CI: [1.7–3.9], $d = 2.0$, $p < 0.0001$); and a higher peak firing rate in the SOL at short than at intermediate by 1.0 pps (95% CI: [0.3–1.7], d = 0.7, p = 0.0036) and long muscle length by 1.9 pps (95% CI: [1.1–2.7], d = 1.3, p < 0.0001), as well as at intermediate than at long muscle length by 0.9 pps (95% CI: [0.2–1.5], d = 0.6, p = 0.0118). Conversely, no difference between muscle lengths was observed in the incomplete SCI group (all p > 0.1500).

*Estimates of synaptic inputs to motoneurons*

There was a significant interaction between group and muscle length for acceleration (p = 0.0354), with no significant interaction between group, muscle length, and muscle (p = 0.2607). Specifically, the control group showed a higher acceleration at short compared with intermediate (d = 0.46, p = 0.0028) and long muscle length (d = 0.51, p = 0.0186), regardless of the muscle. Conversely, no difference between muscle lengths was observed in the incomplete SCI group (all p > 0.9727). There were no significant interactions involving the group factor for attenuation (all p > 0.2343) and brace height (all p > 0.1023).

When considering ΔF, there was a significant interaction between group and muscle length (p = 0.0203), with no significant interaction between group, muscle length, and muscle (p = 0.0765). Specifically, the control group showed a higher ΔF at short compared with long muscle length (d = 0.65, p = 0.0167), regardless of the muscle. Conversely, no difference between muscle lengths was observed in the incomplete SCI group (all p > 0.8960).

When considering normalized ΔF, there was a significant interaction between group, muscle length, and muscle (p < 0.0001). Specifically, the control group showed a higher normalized ΔF in the intermediate compared with the short (by 19%; 95% CI: [3–35], d = 0.79, p < 0.0001) and long (by 29% ; 95% CI: [11–47], d = 1.18, p < 0.0001) muscle lengths; and the incomplete SCI group showed a higher normalized ΔF in the short compared with the intermediate length (by 10% ; 95% CI: [0–19], d = 0.40, p < 0.0406).

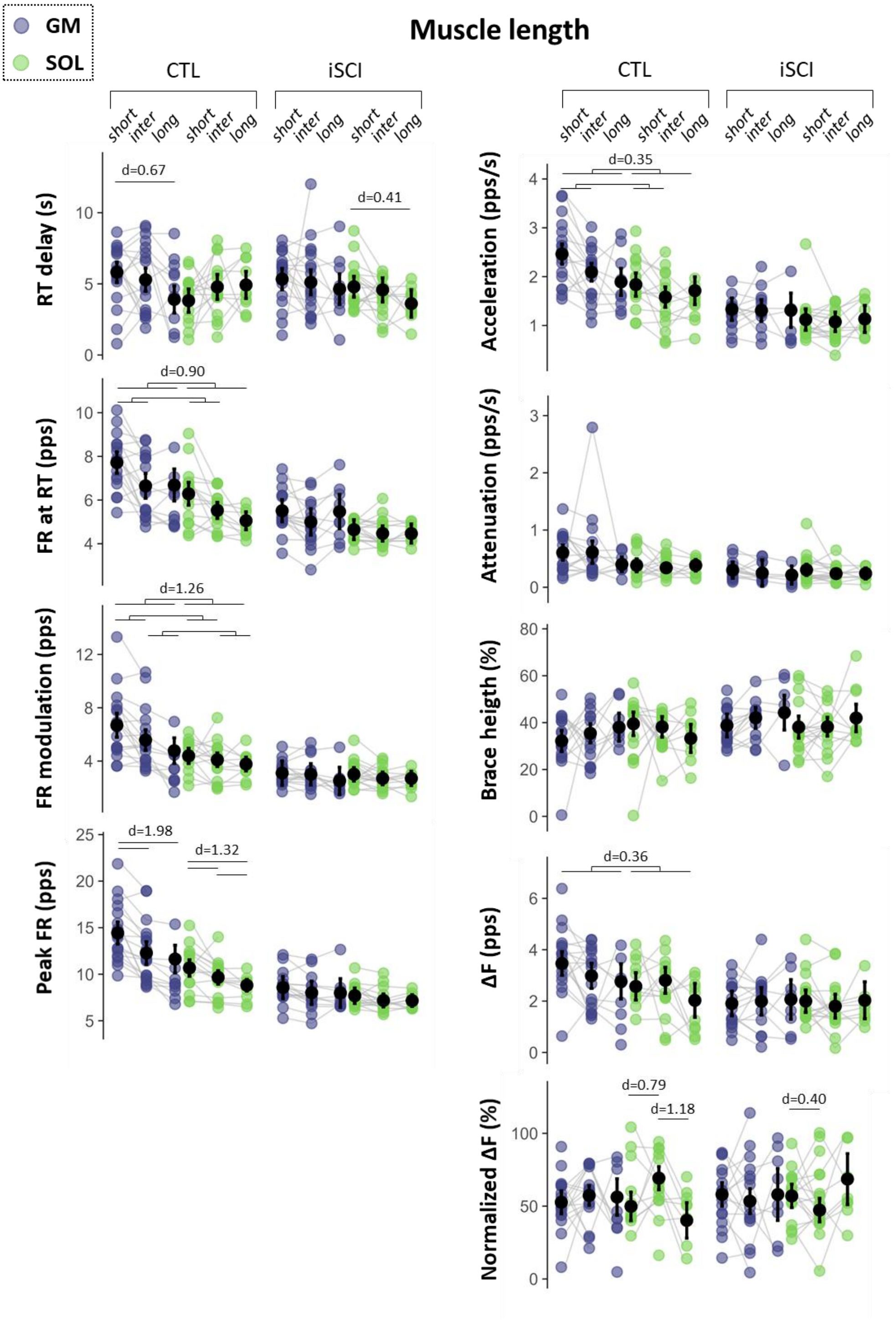


**Supplementary Figure 2. Motoneuron recruitment thresholds, rate coding and estimates of synaptic inputs to motoneurons with changes of muscle lengths, at same absolute torque level.** The incomplete SCI (iSCI) and control (CTL) groups are compared at either the same absolute torque (40% MVT of the short muscle length position). Recruitment threshold (RT), firing rate at recruitment (DR RT), firing rate modulation (FR modulation), and peak firing rate (max DR), acceleration slope, attenuation slope, brace height, ΔF, and normalized ΔF were obtained from the gastrocnemius medialis (GM, purple) and soleus (SOL, green). Colored dots represent participant averages, while black dots and bars indicate the estimated marginal means with 95% confidence intervals predicted by linear mixed-effects models. Cohen's d is reported for significant differences.

### SUPPLEMENTARY ANALYSIS 2 : COMPARING THE EFFECTS OF MUSCLE LENGTH AND VIBRATION IN TRACKED MOTONEURONS

#### Supplementary methods 2

In the core manuscript, to assess how motoneuron behavior changed with changes in muscle length and the application of vibration, the entire set of decomposed motoneurons was used. This approach allows maximization of the number of motoneurons and participants included in the analysis. However, it may lead to comparisons between different motoneurons across conditions. Thus, we complemented our approach by tracking motoneurons across conditions (i.e., changes in ankle angle and application of vibration).

To achieve this, the motor unit filters identified at the short muscle length were applied to the extended and whitened EMG signals of the intermediate muscle length condition (Francic & Holobar, 2021). The resulting spike trains were compared with those initially obtained. A motor unit was considered tracked if at least 30% of the discharge times were shared. Only the motor units tracked across the two muscle lengths were included in the analyses. The same procedure was performed between contractions with and without vibration.

The statistical models used for these comparisons were adapted by adding a random intercept for each motoneuron [e.g., lmer(metric ~ positiongroupmuscle + (positiongroupmuscle|id_participant) + (1|id_participant:id_MU)].

#### Supplementary results 2

*Recruitment threshold and rate coding*

For the following results, the effects of interest are those involving interactions between group and condition (muscle length or vibration), including both group-condition interactions and group-condition-muscle interactions.

For the muscle length condition, there was a significant interaction between group, muscle length and muscle for recruitment threshold (p = 0.0037). Specifically, the control group showed a higher recruitment threshold in the GM at the intermediate than at the short muscle length by 5.5% MVT (95% CI: [1.8–9.1], d = 1.0, p = 0.0037), with no other significant differences (Fig. 3). When considering firing rate at recruitment, there were no significant interactions involving the group factor (all p > 0.1372). Conversely, there was an interaction between group, muscle length and muscle for firing rate modulation (p = 0.0206) and peak firing rate (p = 0.0381). Specifically, the control group showed higher firing rate modulation by 0.86 pps (95% CI: [0.34–1.38], d = 0.80, p = 0.0019) and higher peak firing rate by 1.39 pps (95% CI: [0.79–1.98], d = 1.6, p < 0.0001) in the GM at short compared with the intermediate muscle length, while there was no difference in the SOL or in either muscle in the incomplete SCI group (all p > 0.0730).

For the vibration condition, there were no significant interactions involving the group factor for recruitment threshold (all p > 0.1722). There was an interaction between group and vibration for firing rate at recruitment (p = 0.0212) and for peak firing rate (p = 0.0007), with no significant interaction between group, vibration, and muscle (all p > 0.2815). Specifically, the control group showed a lower firing rate at recruitment (d = 0.75, p = 0.0071) and a lower peak firing rate (d = 1.1, p = 0.0003) in the presence of vibration; whereas the incomplete SCI group showed no significant changes with vibration. There were no significant interactions involving the group factor for rate modulation (all p > 0.1304).

*Estimates of synaptic inputs to motoneurons*

For the muscle length condition, there was no interaction between group and muscle length, nor between group, muscle length, and muscle for acceleration (p > 0.4480), attenuation (p > 0.0756), or brace height (p > 0.0684) (Fig. 4). When considering ΔF, there was an interaction between group, muscle and muscle length (p = 0.0096). Specifically, there were no significant changes in the incomplete SCI group (all p > 0.0783), whereas the control group showed higher ΔF at short muscle length in the GM by 0.43 pps (95% CI: [0.18–0.68], d = 0.62, p = 0.0017) and lower ΔF at short muscle length in the SOL by 0.65 pps (95% CI: [0.17–1.12], d = 0.94, p = 0.0076). By contrast, when considering normalized ΔF, there was no significant interaction between group and muscle length (p = 0.1004), nor between group, muscle length, and muscle (p = 0.6264).

For the vibration condition, there was no interaction between group and vibration, nor between group, muscle length, and muscle for brace height (all p > 0.0791), acceleration (all p > 0.0635), or attenuation (all p > 0.0855). When considering ΔF, there was a significant interaction between group and vibration (p = 0095) with no significant interaction between group, vibration, and muscle (all p > 0.0856). Specifically, there were no significant difference in the incomplete SCI group (p = 0.5155), whereas the control group showed higher ΔF without than with vibration, regardless of the muscle (d = 0.74, p = 0.0060). By contrast, when considering normalized ΔF, there was no significant interaction was found between group and vibration (p = 0. 7497), nor between group, vibration, and muscle (p = 0.2307).

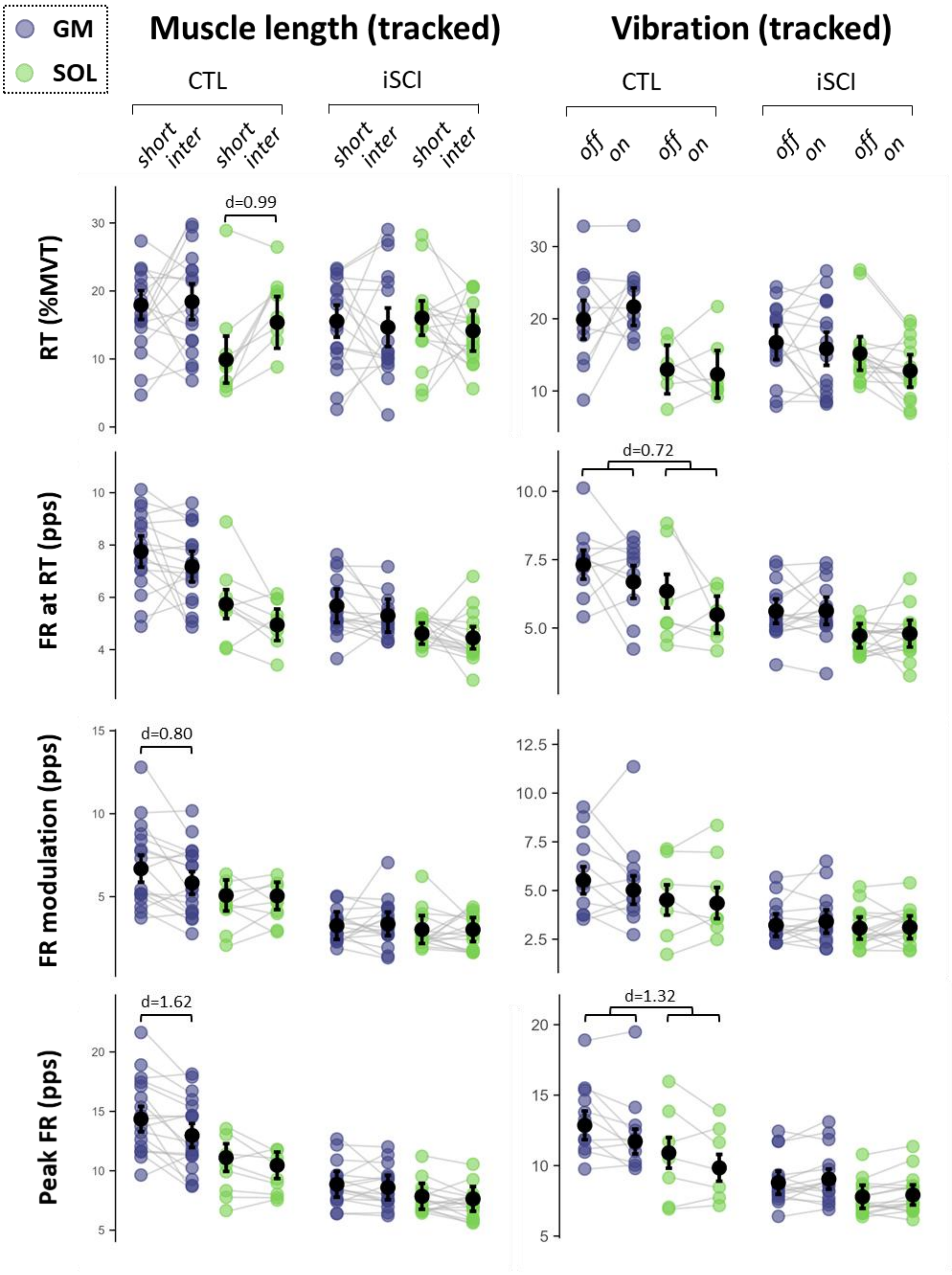


**Supplementary Figure 3. Effect of muscle length and vibration in motoneuron recruitment threshold and rate coding, with tracked motoneurons across conditions.** The modulation of motoneuron rate coding and recruitment threshold between the incomplete SCI (iSCI) and control (CTL) groups are compared in response to changes in muscle length at the same relative torque (40% MVT), and the application of tendon vibration to antagonist muscles, at short muscle length (20° of plantarflexion). Recruitment threshold (RT), firing rate at recruitment (DR RT), firing rate modulation (FR modulation), and peak firing rate (peak FR), were obtained from the gastrocnemius medialis (GM, purple) and soleus (SOL, green). Colored dots represent participant averages, while black dots and bars indicate the estimated marginal means with 95% confidence intervals predicted by linear mixed-effects models. Cohen's d is reported for significant results ($p < 0.05$).

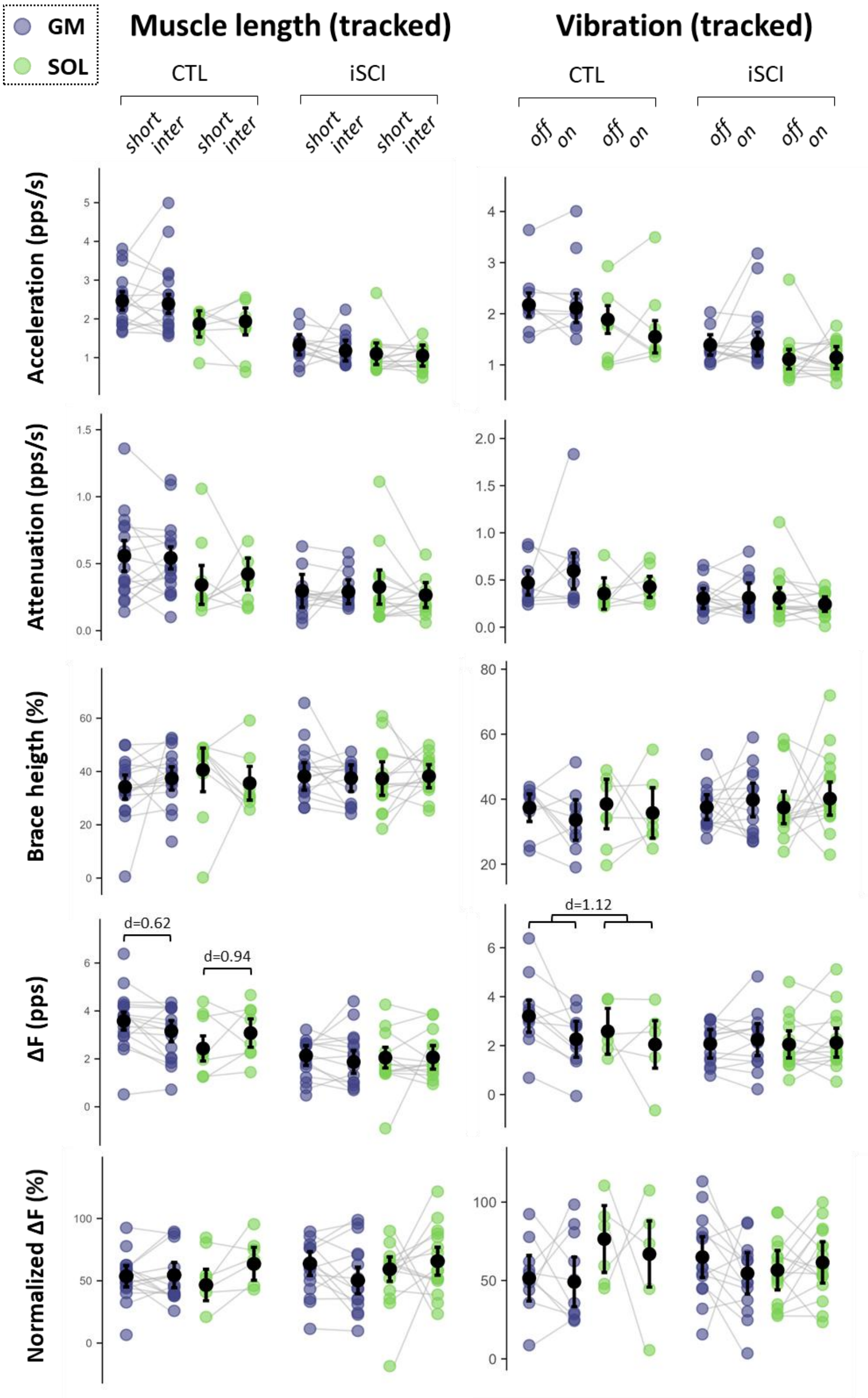


**Supplementary Figure 4. Effect of muscle length and vibration in estimates of synaptic inputs to motoneurons, with tracked motoneurons across conditions.** The modulation of motoneuron estimates of synaptic inputs to motoneurons between the incomplete SCI (iSCI) and control (CTL) groups are compared in response to changes in muscle length at the same relative torque (40% MVT), and the application of tendon vibration to antagonist muscles, at short muscle length (20° of plantarflexion). The acceleration slope, attenuation slope, brace height, ΔF, and normalized ΔF were obtained from the gastrocnemius medialis (GM, purple) and soleus (SOL, green). Colored dots represent participant averages, while black dots and bars indicate the estimated marginal means with 95% confidence intervals predicted by linear mixed-effects models. Cohen’s d is reported for significant results ($p < 0.05$).
